# Surveying the sweetpotato rhizosphere, endophyte, and surrounding soil microbiomes at two North Carolina farms reveals underpinnings of sweetpotato microbiome community assembly

**DOI:** 10.1101/758359

**Authors:** C Pepe-Ranney, C Keyser, J Trimble, B Bissinger

## Abstract

Farmers grow sweetpotatoes worldwide and some sub-Saharan African and Asian diets include sweetpotato as a staple, yet the sweetpotato microbiome is conspicuously less studied relative to crops such as maize, soybean, and wheat. Studying sweetpotato microbiome ecology may reveal paths to engineer the microbiome to improve sweetpotato yield, and/or combat sweetpotato pests and diseases. We sampled sweetpotatoes and surrounding soil from two North Carolina farms. We took samples from sweetpotato fields under two different land management regimes, conventional and organic, and collected two sweetpotato cultivars, ‘Beauregard’ and ‘Covington’. By comparing SSU rRNA gene amplicon sequence profiles from sweetpotato storage root skin, rhizosphere, and surrounding soil we found the skin microbiome possessed the least composition heterogeneity among samples and lowest alpha-diversity and was significantly nested by the rhizosphere in amplicon sequence variant (ASV) membership. Many ASVs were specific to a single field and/or only found in either the skin, rhizosphere, or surrounding soil. Notably, sweetpotato skin enriched for *Planctomycetaceae* in relative abundance at both farms. This study elucidates underpinnings of sweetpotato microbiome community assembly, quantifies microbiome composition variance within a single farm, and reveals microorganisms associated with sweetpotato skin that belong to common but uncultured soil phylotypes.

## Introduction

The microbiome influences plant health and development, playing a vital role at all stages of plant growth. Microorganisms affect plant tolerance to drought (Fitzpatrick et al. 2018) and disease (Bakker et al. 2018), enhance the plant’s ability to acquire nutrients (Friesen et al. 2011), and impact yield (Busby et al. 2017). As such, engineering the crop microbiome may be of considerable economic value (Toju et al. 2018). Many studies have investigated the microbiomes of crops commonly grown in the United States such as maize (Emmett et al. 2017; Bouffaud et al. 2014; Peiffer and Ley 2013; Niu et al. 2017; Peiffer et al. 2013), soybean (Wang et al. 2017; Rascovan et al. 2016; Zhang et al. 2018; Hamid et al. 2017), and wheat (Mahoney, Yin, and Hulbert 2017; Yin et al. 2013, 2017; Rascovan et al. 2016; Gdanetz and Trail 2017; Mavrodi et al. 2018; Donn et al. 2015). In contrast, few studies have investigated the microbiome of sweetpotato.

Sweetpotato (*Ipomoea batatas*) is an important food crop worldwide (Kays 2004). In the United States, sweetpotato production has varied over the last century, primarily influenced by the economy, but it continues to be an important crop (Smith et al. 2009). In Asia and sub-Saharan Africa (SSA) sweetpotato provides crucial nutrients (Low et al. 2017, 2009). Orange-fleshed varieties of sweetpotato contain beta-carotene which is converted to Vitamin A, a nutrient commonly deficient in SSA diets (Low et al. 2017). Presently, China produces the most sweetpotatoes by country (>24 million tons on average from 2012-2014, Low et al. (2017)). SSA has more land under sweetpotato cultivation than China, and SSA land use for sweetpotato cultivation is expanding faster than the land area under maize cultivation (Low et al. 2017, 2009).

Given the worldwide nutritional value of sweetpotatoes, it is desirable to explore the potential of the microbiome to promote sweetpotato yield, drought tolerance, and to manage sweetpotato pests and disease. To date, culture-independent and -dependent techniques have elucidated the impact of some sweetpotato genotypes on sweetpotato rhizosphere microbiome composition (Marques et al. 2014) and produced isolated microorganisms in culture from the sweetpotato endophyte community (Marques et al. 2015). Rhizosphere versus surrounding soil contained different microbial community composition in three sweetpotato genotypes (Marques et al. 2014). In addition, sweetpotato genotype correlated with rhizosphere bacterial community composition but represented much less variance in microbial community composition than the variance represented by sweetpotato compartment (i.e. bulk soil versus rhizosphere sample types). Isolates from the sweetpotato endophytic compartment included seventeen distinct genera from 93 individual isolates. Over half of the isolates, 51 of 93, belonged to the genus *Bacillus*. Twenty-five of the 93 isolates were tested for antagonistic activity against *Plenodomus destruens*, a sweetpotato fungal pathogen, of which four *Bacillus* isolates appeared to kill *P. destruens* in an agar overlay assay (Marques et al. 2015).

Complicating meta-analyses of plant microbiome research, methods for studying microbiomes improve rapidly with advances in DNA sequencing technology. Microbiome studies for a given plant should be renewed with the latest technology to keep the collective understanding of microbiome biology current. Notable recent advancements include: the ability to distribute small subunit (SSU) rRNA gene amplicon sequence reads into high resolution ecological units, such as “amplicon sequence variants” (ASVs) (Callahan et al. 2016), “oligotypes”, and minimum entropy decomposition “nodes” (Eren et al. 2013, 2015); the adoption of statistical methods by microbial ecologists for studying compositional data (Morton et al. 2017; Silverman et al. 2017; Quinn et al. 2018); and improved techniques for estimating variance in alpha-diversity indices (Willis and Martin 2018). The complexity of the soil microbiome makes it difficult to interpret the influence of the microbiome on agroecosystem productivity, yet we can interrogate microbial communities at ever-increasing scale and resolution. We can also place results in the context of other studies through direct, data-driven comparisons as standards, practices, and data curation/availability improve (Thompson et al. 2017).

Elucidating rules of community assembly for the agricultural phytobiome has been identified as a key research need to make use of the microbiome to improve agricultural productivity (Busby et al. 2017). Models for plant-associated microbiome community assembly have been tested for many crops (e.g. Edwards et al. 2015) and other plants (Bulgarelli et al. 2013) although no studies have focused on sweetpotato-associated microbiome assembly rules. To expand upon the small body of sweetpotato microbiome research, we profiled the sweetpotato microbiome across multiple fields on a single farm and in one field at a nearby working farm in North Carolina. Our study quantifies the relative changes in microbiome composition across sweetpotato storage root skin, rhizosphere and surrounding soil. To provide a broad context for interpretation and better understand how our results might be generalized, we collected sweetpotatoes across several fields under different land management regimes (conventional versus organic), two different sweetpotato cultivars, ‘Beauregard’ and ‘Covington’, and collected sweetpotatoes at two working North Carolina sweetpotato farms.

## Results

We visited two working North Carolina sweetpotato farms, Burch Farm and Sharp Farm, and collected samples from three sweetpotato fields at Burch Farm and one field at Sharp Farm. At Burch Farm, the three field locations included four distinct sweetpotato plots: two *conventionally* managed plots at two distinct field locations where each plot had one sweetpotato cultivar, ‘Beauregard’ or ‘Covington’, and two *organically* managed plots at one field location where each plot contained one sweetpotato cultivar, again ‘Beauregard’ or ‘Covington’ (Figure 1, Table 1). We collected from one field at Sharp Farm growing the ‘Covington’ cultivar under conventional management practices. At each sampling location, we collected crop-adjacent soil and a sweetpotato storage root from ∼15 sites (Table 1). Sweetpotato storage root samples were divided into two subsamples: skin and rhizosphere. The ‘skin’ samples served as a proxy for the sweetpotato endophytic compartment. For each sweetpotato skin, rhizosphere, and surrounding soil sample we produced and sequenced on average more than 14,000 high-quality V4 SSU rRNA gene amplicon sequences (Table 1, Methods).

**Figure 1.**
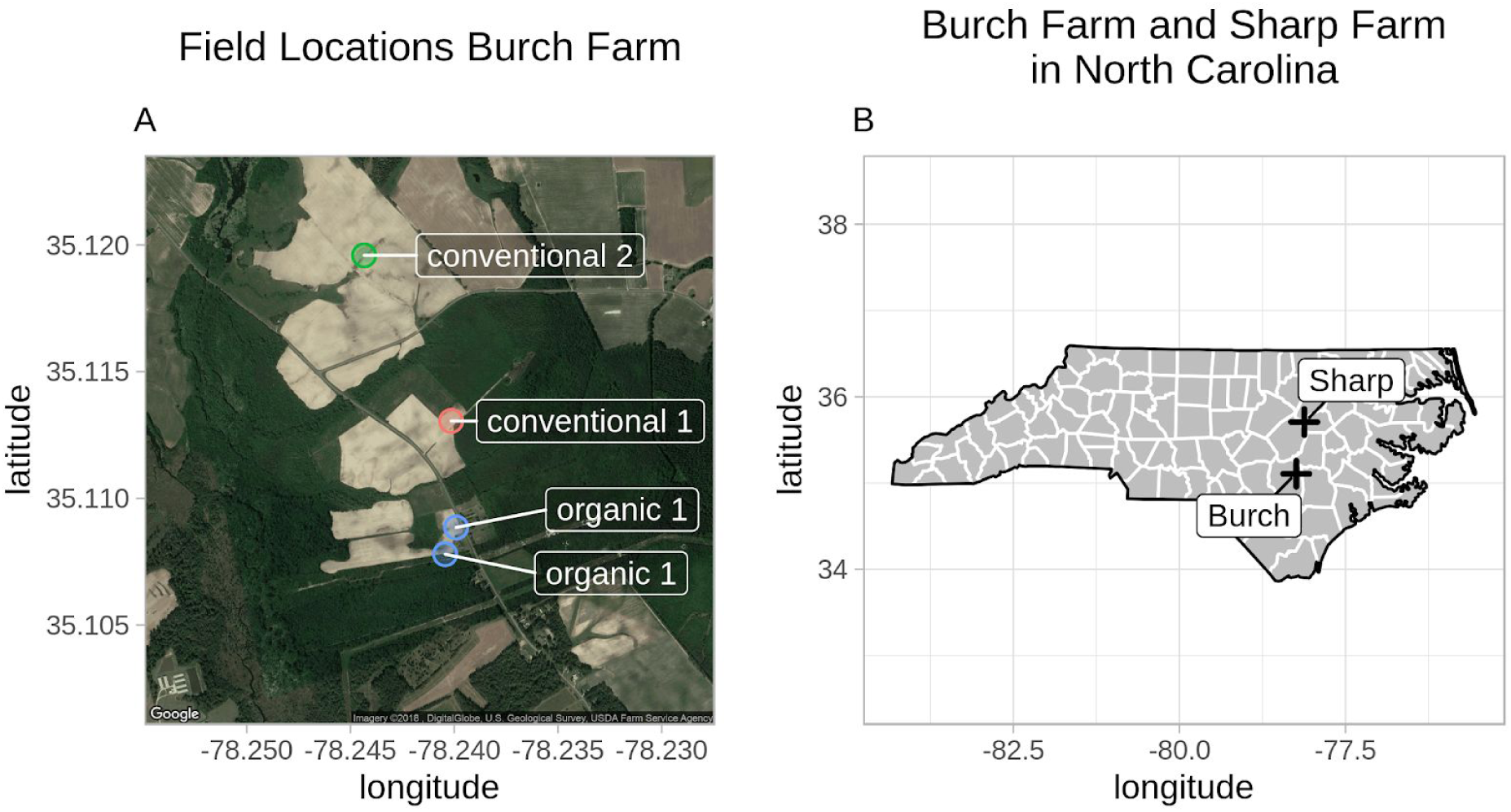
Farm and field locations. Field locations within Burch Farm (**Panel A**) and the locations of Burch and Sharp Farms in North Carolina **(Panel B)**. The ‘organic 1’ field location had two plots: one with the ‘Beauregard’ cultivar and one with ‘Covington’.

**Table 1.** Sample information.

| Field location | Cultivar* | Compartment* | Number of samples† | mean obs.‡ | std. dev. obs.§ |
| --- | --- | --- | --- | --- | --- |
| **Burch Farm** |  |  |  |  |  |
| conventional 1 | Covington | rhizosphere | 15 | 18,697 | 2,731 |
|  | Covington | skin | 14 | 12,348 | 2,630 |
|  | Covington | soil | 15 | 16,558 | 2,974 |
| conventional 2 | Beauregard | rhizosphere | 15 | 13,741 | 1,303 |
|  | Beauregard | skin | 11 | 7,251 | 1,435 |
|  | Beauregard | soil | 14 | 12,058 | 1,275 |
| organic 1 | Beauregard | rhizosphere | 15 | 21,182 | 3,909 |
|  | Covington | rhizosphere | 13 | 13,502 | 1,415 |
|  | Beauregard | skin | 14 | 15,960 | 4,179 |
|  | Covington | skin | 13 | 10,706 | 1,749 |
|  | Beauregard | soil | 12 | 15,796 | 3,040 |
|  | Covington | soil | 12 | 10,950 | 1,446 |
| **Sharp Farm** |  |  |  |  |  |
| conventional 3 | Covington | rhizosphere | 14 | 17,745 | 1,771 |
|  | Covington | skin | 13 | 11,700 | 2,516 |
|  | Covington | soil | 14 | 13,496 | 2,506 |

\* Colors correspond to the legends in Figure 2.
† Number of samples in this study after DNA sequence quality control
‡ Mean observations (i.e. ASV observations) produced from samples
§ Standard deviation of observations produced from samples

### Microbial community composition clustered by compartment and field

Sweetpotato skin, rhizosphere, and crop-adjacent soil microbial communities differed from one another in phylogenetic composition (Figure 2, PERMANOVA p-value 0.001). Additionally, microbial community composition varied across fields within Burch Farm (Figure 2, PERMANOVA p-value 0.001). Although the field layout at Burch Farm did not enable us to disentangle the variance represented by field location versus land management regime (organic or conventional) (Figure 1), we note that fields within the same farm under different land management regimes demonstrated striking differences in community composition (Figure 2). Sweetpotato compartment microbial communities from Sharp Farm were similar to communities of the same compartment from Burch Farm (Figure 2) -- that is, Sharp Farm rhizosphere samples clustered with Burch Farm rhizosphere samples, crop-adjacent soil with crop-adjacent soil, and sweetpotato skin with sweetpotato skin (see NMDS ordination of Bray-Curtis distances between samples Figure 2).

**Figure 2.**
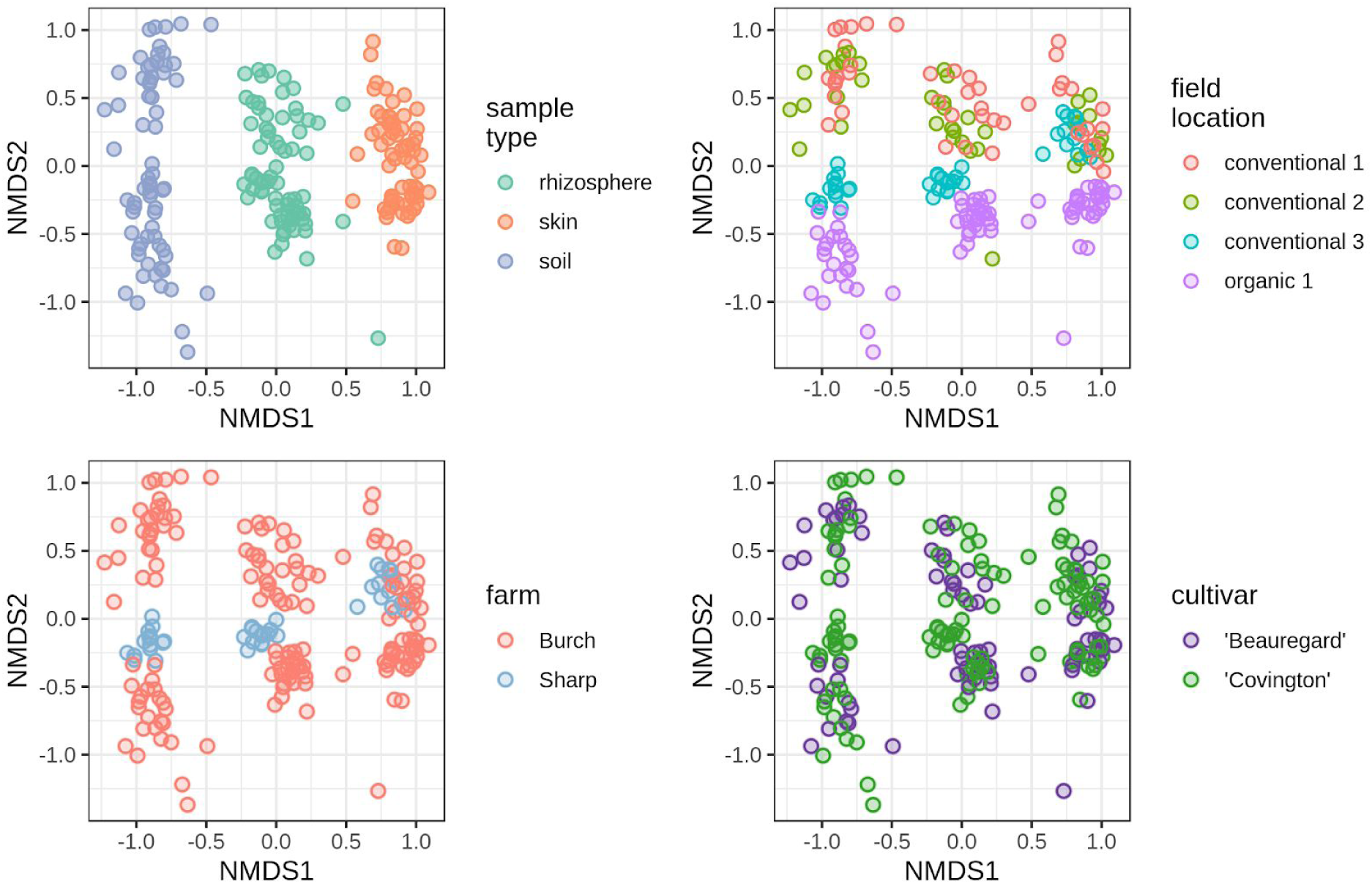
NMDS ordination of Bray-Curtis distances between samples. Non-metric multidimensional scaling (NMDS) ordination of Bray-Curtis distances between samples based on ASV content. Every panel depicts the same ordination. Points are samples. Point colors for each panel correspond to a different sample category as indicated in the legends.

### Broad and narrow resolution phylogenetic differences contributed to community composition changes across sweetpotato compartments

We found phylogenetic balances that separated samples by compartment within each farm by fitting a penalized multinomial logistic regression model of sweetpotato microbiome compartment with phylogenetic ILR transform values (i.e. ‘phylogenetic balances’) (see Methods) (Silverman et al. 2017; Friedman, Hastie, and Tibshirani 2009). Briefly, a ‘balance’ is the ratio of descendent taxa abundances on either side of a dendrogram node. The dendrogram can be any bifurcating representation of taxa relationships. Possible topology bases from which to calculate balances include phylogenies and hierarchical clusterings of taxa by abundances across samples (Morton et al. 2017). Balances are preferred to relative abundances (i.e. proportions) as analyses based on simple taxa proportions lead to spurious results (Gloor et al. 2017; Friedman and Alm 2012; Quinn et al. 2018). A ***‘phylogenetic*** balance’ is a transformation of microbial community taxa counts into balance values based on a phylogenetic tree that incorporates phylogenetic information such as branch lengths (see Silverman et al. (2017) for full derivation).

### Burch Farm

Balances from varying depths in the taxa phylogeny separated samples by compartment (Figure 3, Supplementary Figure 1). For example, taxa abundances in the *Acidobacteria* versus abundances in the *Gemmatimonadetes*, *Firmicutes*, *Actinobacteria*, *Proteobacteria,* and other phyla separated rhizosphere and skin samples from surrounding soil (balance ‘n5’, Figure 3, Supplementary Figure 1). A number of balances with members in the *Alphaproteobacteria* and adjacent classes also separated skin and rhizosphere samples from soil (balances ‘n138’, ‘n140’, ‘n146’, ‘n149’, ‘n153’, ‘n162’, and ‘n164’). For example, balance ‘n153’’ strongly separated skin and rhizosphere from soil samples. The sister clades that made up balance ‘n153’ included a clade that was predominantly *Alphaproteobacteria* and a smaller clade consisting of ASVs in the *Sphingomonadaceae.* One balance, ‘n138’, had one clade with members in the *Alphaproteobacteria*. The other clade that made up balance ‘n138’ included two members not closely related to any cultured microorganism but that shared 100% sequence identity with environmental 16S rRNA gene clone sequences from a number of culture-independent studies. Matches to one member of the non-*Alphaproteobacteria* clade of balance ‘n138’ included 16S rRNA gene sequences from unpublished studies of winery wastewater (Genbank accession MF654612), anaerobic digesters (accession MG854374) and a published study of a flower shop daisy rhizosphere ((Tabei and Ueno 2010), accession AB511015). The other member of the ‘n138’ non-*Alphaproteobacteria* sister clade shared 100% sequence identity with 16S rRNA gene sequences recovered in an unpublished study of cropping systems in China (accession KR836688) and a published study of *Bdellovibrio*-and-like organisms ((Davidov and Jurkevitch 2004), accession AY294213). Balances ‘n140’, ‘n146’, and ‘n149’ varied similarly across compartment and each consisted of a one small clade (3-6 members) that either generally increased or remained constant in relative abundance with distance from the sweetpotato (i.e. from skin to rhizosphere to adjacent soil) and a large clade (75 to 82 members) that generally decreased in abundance with distance from the sweetpotato (Supplementary Figure 1).

**Figure 3.**
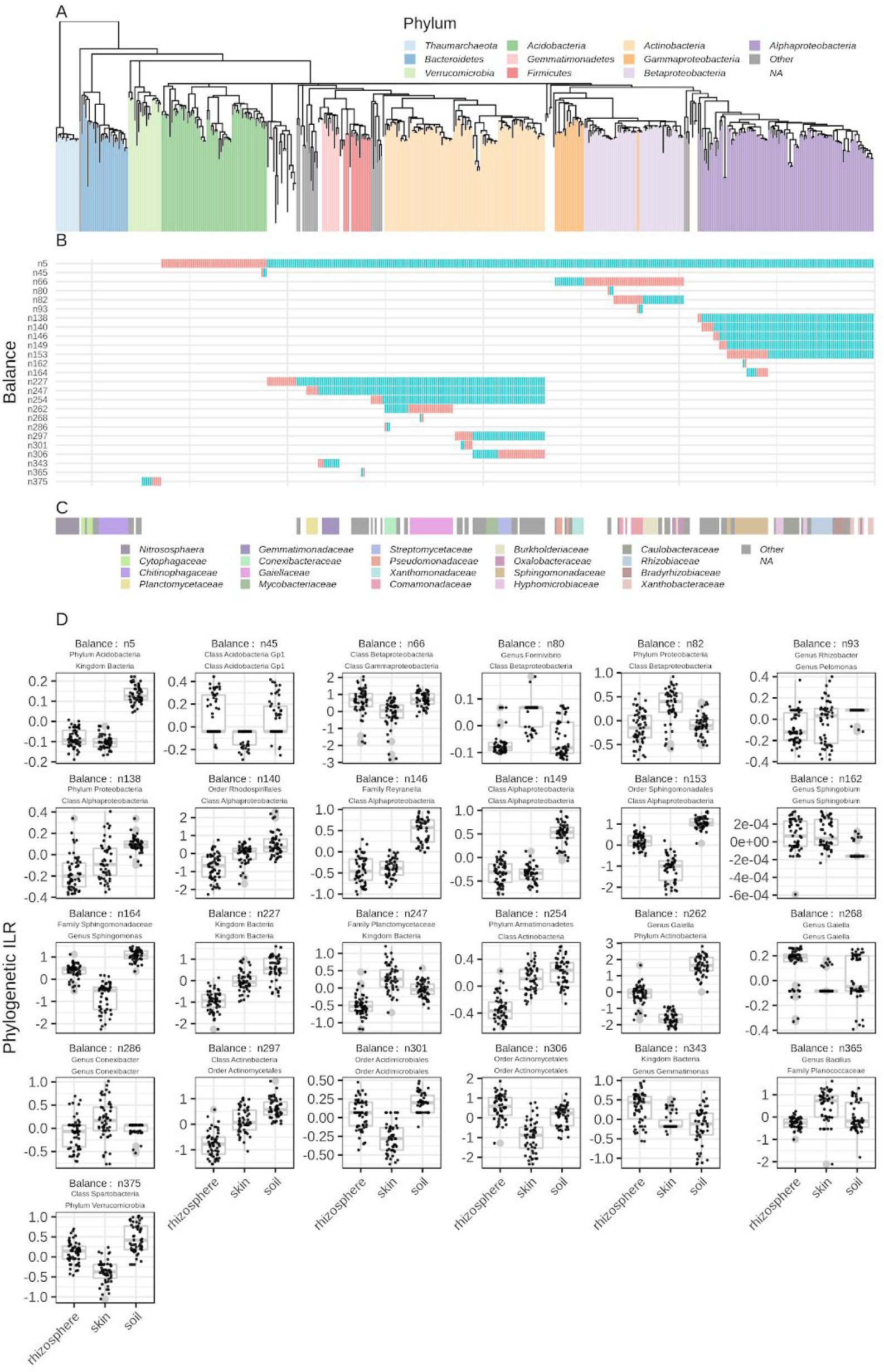
Phylogenetic balances that separated samples by compartments at Burch Farm. Position and phylogenetic ILR (Silverman et al. 2017) values for balances found to separate samples by compartment at **Burch Farm**. **Panel A** shows the phylogenetic relationship of ASVs used in the phylogenetic balance analysis (see Methods). **Panel B** shows the positioning in the phylogeny of each balance. Colors in the second panel denote which side (i.e. sister clade) of the balance was used as the denominator (blue) or numerator (red) when calculating the phylogenetic ILR for each balance. **Panel C** shows the position of Family-level taxonomic groups in the tree. **Panel D** shows the balance ILR values grouped by compartment for each balance highlighted in the second panel. Taxa names in the title of each **Panel D** subplot are the highest resolution, 95% consensus taxonomic affiliation for the sister clades that makes up the balance. The consensus taxonomy of the sister clade used as the numerator to calculate the balance phylogenetic ILR is listed first followed by the denominator clade. Boxplot boxes extend from the 25th to the 75th percentile and include a horizontal line indicating the median. Whiskers extend to the largest value that is no greater than 1.5 times the interquartile range from the 25th (bottom of box) or 75th (top of box) percentile. Points beyond the whiskers are plotted individually.

Eight more balances of moderate phylogenetic breadth distinguished sites by compartment (balances ‘n66’, ‘n82’, ‘n227’, ‘n247’, ‘n254’, ‘n262’, ‘n297’, and ‘n306’, Figure 3, Supplementary Figure 1). The balance ‘n66’ consisting of the split between *Gammaproteobacteria* and *Betaproteobacteria* classes separated the skin from rhizosphere and soil samples. Balance ‘n82’ marked a split within the *Betaproteobacteria* between a clade of predominantly *Comamonadaceae* members and clade with a combination of *Oxalobacteraceae* and *Burkholderiaceae* members. Balance ‘n247’ also separated sweetpotato skin from rhizosphere and soil samples. The sister clades that make up balance ‘n247’ included a clade comprised of members from the *Planctomycetaceae* Family and a clade of *Gemmatimonadetes*, *Firmicutes*, *Actinobacteria* and other phyla. Balances ‘n262’ and ‘n306’ both had the lowest values in skin samples. Balance n262 consisted of a clade with members in the *Gaiella* that decreased in relative abundance with proximity to the sweetpotato and a clade with predominantly *Conexibacter* and *Solirubrobacter* members that increased in relative abundance in the skin versus the rhizosphere and surrounding soil. Balance ‘n306’ was less clearly defined by taxonomic membership. Notable members of one ‘n306’ sister clade included the *Mycobacterium* and *Nocardoides* genera while the other clade in the ‘n266’ balance included members of *Streptomyces* and *Arthrobacter*. Balance ‘n297’ had clades that separated members consistent with the taxonomic rank of order one clade being entirely *Acidimicrobiales* and the other *Actinomycetales*. The *Actinomycetales* clade was greatest in relative abundance in the rhizosphere.

Seven narrow balances with three or less ASV members also separated samples by compartment (balances ‘n45’, ‘n80’, ‘n93’, ‘n162’, ‘n268’, ‘n286’, and ‘n365’, Figure 3, Supplementary Figure 1). These five balances had members in six different phyla. Balance ‘n45’ of strongly separated samples by compartment. This balance included three ASVs in the *Acidobacteria*. One sister clade in balance ‘n45’ was a singleton found in 27 and 25 rhizosphere and soil samples, respectively, but not found in any skin samples. The ASV member of this singleton clade shared 95% SSU rRNA gene sequence identity with *Acidicapsa ligni* strain WH120 (Kulichevskaya et al. 2012) isolated from peat and decaying wood and *Acidobacterium ailaaui* strain PMMR2 (Myers and King 2016) isolated from a geothermally heated Hawaiian microbial mat. One or both sister clades that made up the balances ‘n93’, ‘n162’, and ‘n286’ increased in relative abundance in sweetpotato skin relative to surrounding soil. These balances may identify microbes adapted to the sweetpotato endophytic compartment across three separate classes/phyla, *Betaproteobacteria*, *Alphaproteobacteria*, and *Actinobacteria*.

Notably, phylogenetic ILR values for balance ‘n375’, with members entirely in the *Verrucomicrobia,* decreased from soil to rhizosphere to sweetpotato skin samples (Figure 3, Supplementary Figure 1). This was largely due to decreasing relative abundance of one sister clade in ‘n375’ with proximity to the sweetpotato (i.e. from soil to sweetpotato skin).

### Sharp Farm

We also identified balances that separated samples on the basis of compartment affiliation at Sharp Farm (Supplementary Figure 2, Supplementary Figure 3). Similar to Burch Farm, at Sharp Farm balances ‘n5’ and ‘n247’ -- consisting of sister clades that marked the taxonomic split of *Acidobacteria* and *Planctomycetaceae* from several phyla (see above) for ‘n5’ and ‘n247’, respectively -- separated samples by compartment. Both balance ‘n5’ and ‘n247’ displayed similar trends across compartments whether at Sharp or Burch Farms. Balance ‘n138’ also separated samples by compartment at both farms. The general trends in relative abundance of the balance ‘n138’ sister clades were consistent at each farm.

Four other balances of moderate phylogenetic breadth (balances ‘n194’, ‘n253’, ‘n261’, and ‘n307’) separated samples by compartment (Supplementary Figure 2, Supplementary Figure 3). For both balances ‘n194’ and ‘n153’ balance values were lowest in the surrounding soil. One sister clade of balance ‘n194’ had members predominantly of the *Rhizobiaceae* family while the other clade included members in the *Methylobacteriaceae*, *Bradyrhizobiaceae*, and *Xanthobacteraceae*. The ‘n194’ balance value trends were consistent for Burch Farm and Sharp Farm. Balance ‘n253’ had one clade of predominantly *Actinobacteria* and *Armatimonadetes* members and one clade of *Firmicutes*, *Gemmatimonadetes* members. Balance ‘n253’ had the highest values in rhizosphere and skin samples at Sharp Farm but not at Burch Farm. Unlike at Sharp Farm, the *Actinobacteria*/*Armatimonadetes* clade had greater relative abundances than the *Firmicutes*/*Gemmatimonadetes* clade at Burch Farm. Balances ‘n261’ and ‘n307’ both had members entirely in the *Actinobacteria* and had highest values in skin and soil versus the rhizosphere samples. The two sister clades of balance ‘n261’ were largely consistent with the taxonomic split of orders *Gaiellales* and *Solirubrobacterales* with the orders *Acidimicrobiales* and *Actinomycetales*. Balance ‘n307’ had one clade with most members in the *Streptomyces* genus and one clade with members all in the *Actinomycetales* order.

There were six more narrow balances (‘n34’, ‘n156’, ‘n166’, ‘n335’, ‘n362’, and ‘n409’) that separated samples by compartment (Supplementary Figure 2, Supplementary Figure 3). Balance ‘n34’ strongly separated sweetpotato skin from rhizosphere and soil samples and consisted of a singleton clade whose member ASV was in the *Acidobacteria* Group 1 order and a sister clade of taxa in a number of *Acidobacteria* orders in including *Acidobacteria Group 1*, *Granulicella*, and *Terriglobus*. Balance ‘n34’ did not display similar trends across compartments at Burch Farm. Balance ‘n156’ consisted of a singleton clade that was not found in any Sharp Farm sample and another clade with four members of the *Sphingomonadales* order that generally increased in relative abundance with proximity to the sweetpotato. Both ‘n156’ clades were observed at Burch Farm. Balance ‘n166’ had only two members and strongly differentiated skin from rhizosphere samples. Each member ASV of balance ‘n166’ was in the *Sphingomonadaceae* family. Notably, balance ‘n335’ had a singleton clade with an ASV member only observed at Sharp Farm in skin or rhizosphere samples. This ASV member was also more prevalent in the skin and rhizosphere at Burch Farm although it was observed in the Burch Farm crop-adjacent soil. Balances ‘n362’ and ‘n409’ were made of clades not observed in the skin samples at Sharp Farm. Balance ‘n362’ is comprised of two singletons. One singleton ASV from ‘n362’ was rarely found in the soil or skin at either Burch or Sharp Farm. This ASV shares 100% sequence identity with two isolates from the *Paenibacillus* (*Paenibacillus taohuashanense* strain gs65 isolated from rhizosphere soil (Xie et al. 2012) and *Paenibacillus wynnii* isolated from Alexander Island, Antarctica (Rodríguez-Díaz et al. 2005)).

### High-resolution ASVs displayed strikingly low distribution entropy across compartments and fields

From the set of 14,148 ASVs that passed quality control and were observed at least 10 times, 8,343 ASVs (8.2% of observations from ASVs observed 10 or more times) were found in only one compartment. There were 9,435 ASVs (12.8% of observations from ASVs observed 10 or more times) found at only one field location, and 7,340 ASVs (6.1% of observations from ASVs observed at least 10 times) were found in only one compartment at one field location. We calculated the field/compartment distribution entropy for each of the above 14,148 ASVs (see Methods) and for context contrasted entropy at the ASV-level with the field/compartment distribution entropy at the taxonomic rank of genus. Genera clearly displayed higher field location plus compartment distribution entropy than ASVs (Figure 4). That is to say, taxa at the genus-level are more widespread than ASVs. This is expected as the genus-level is a much broader category in phylogenetic breadth than the ASV-level, however, the effect on agroecosystem productivity of compartment/field specific ASV content is unclear.

**Figure 4.**
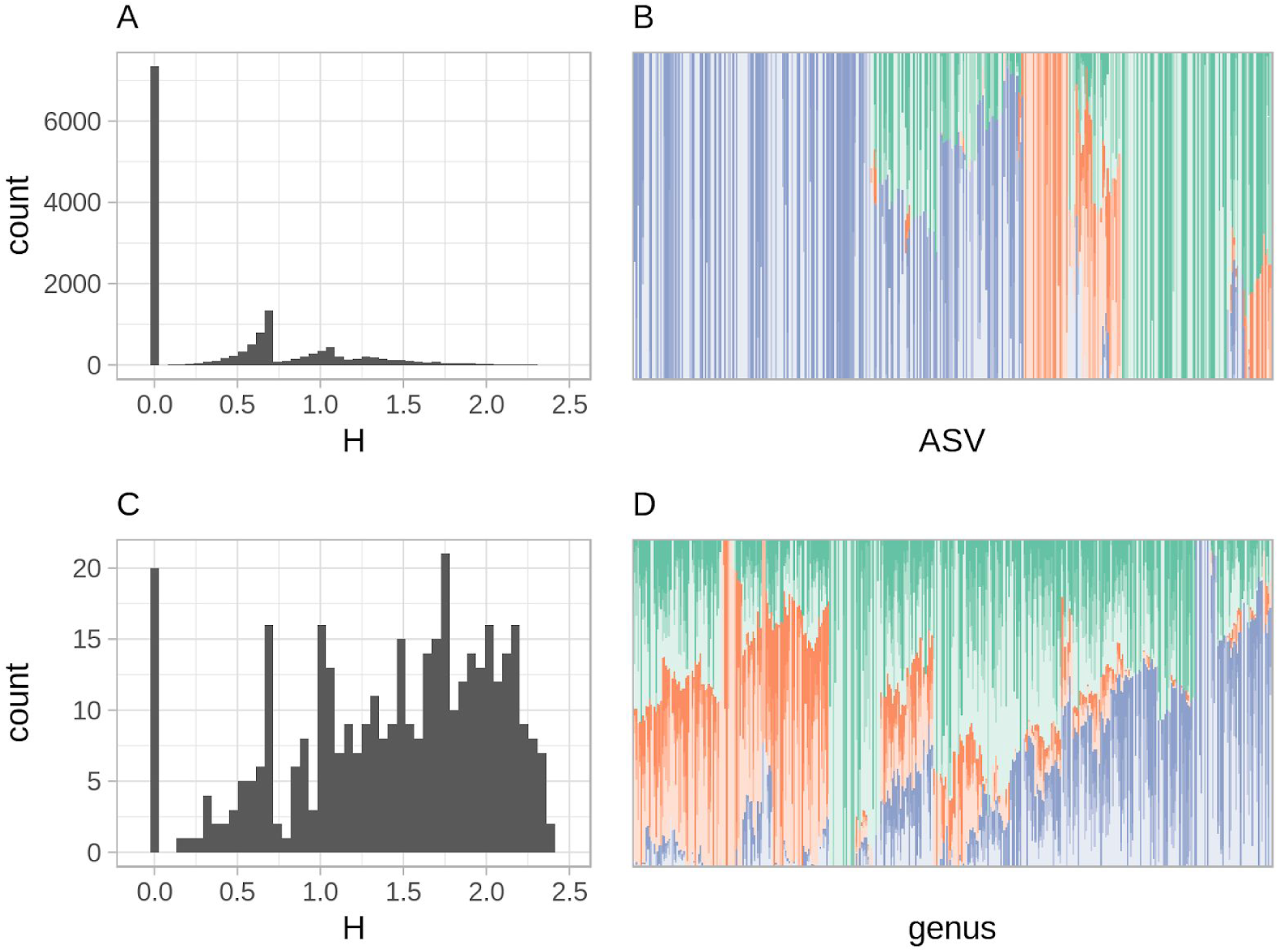
Distribution entropy of ASVs and genera across field-compartment combinations. **Panels A and B** display sample distribution entropy (see Methods) at the ASV-level across field plus compartment categories. **Panels C and D** display sample distribution entropy at the genus-level. **Panels A and C** show the histogram of distribution entropies for ASVs with at least ten observations (**A**), and all genera (**C**). **Panels B and D** show the compartment/field location distribution of ASVs (**B)** or genera (**D**). Colors denote the compartment and are consistent with the legend in Figure 2. Each field location is shaded differently (i.e. ‘conventional 1’, ‘conventional 2’, ‘conventional 3’, and ‘organic 1’ are shaded from **dark** to **light**). All 390 genera (**D**) or a random sample of 390 ASVs (**B**) are shown to keep the number of taxa consistent across each panel. As an example, a solid vertical bar indicates the taxon (ASV or genus) was found in only one field and one compartment. Bars depict the compartment/field values for all observations of the given ASV or genus.

Many compartment-specific ASVs (5,375 of 8,343) were not classified at the genus-level using our taxonomic classification criteria. Of the 5,375 ASVs not classified at the genus-level, 1,159 were also not classified at the taxonomic rank of phylum. To assess the global prevalence of these taxonomically undescribed ASVs, we checked for their presence in a subset of the Earth Microbiome Project (EMP) (Thompson et al. 2017) representing soil-specific data (see Methods, Supplementary Figure 4). We found 782 of the 1,159 compartment-specific OTUs had a match of at least 99% shared sequence identity to an ASV from the EMP. A greater fraction of the compartment-specific ASVs, 4,065 of 5,375, that could not be affiliated with any genus matched at least one ASV in the EMP dataset with equal to or greater than 99% sequence identity. Of the 5,375 compartment-specific ASVs not affiliated with a genus, 2,805 matched another ASV from this study data at greater than or equal to 99% sequence identity.

The five most common genus-annotations for compartment-specific ASVs that were annotated at the genus-level were *Gemmatimonas*, *Gemmata*, *Aquicella*, *Conexibacter*, and *Zavarzinella* (Supplementary Figure 5). The fifteen most abundant compartment-specific ASVs by total sequence count were often found in more than one field and all were found in more than one sample (Table 2). We also note that many of the most abundant compartment-specific ASVs could be annotated taxonomically at better than the phylum-level (Table 2). All except one of the top fifteen compartment-specific ASVs by total sequence count matched at least one ASV in the EMP with 100% sequence identity. The one compartment-specific ASV (‘`8d5236ff`’, Table 2) that did not match an EMP ASV with at least 100% sequence identity was specific to the rhizosphere compartment and is most closely related to *Solimonas aquatica* strain NAA16 among cultured isolates (97% shared V4-region SSU rRNA gene sequence identity, accession NR_108189). One of the fifteen most numerous compartment-specific ASVs (‘`c194bf20`’, Table 2), is affiliated with the candidate division WPS-2 (Writtenberg Polluted Soil) (Nogales et al. 2001) and matched an EMP ASV with 100% sequence identity that was observed in 39 EMP samples. Of the 39 EMP samples, 38 appear to be agricultural soil with EMP ‘`env_feature`’ values of ‘agricultural feature’ (17 samples), ‘cultivated habitat’ (8 samples), ‘agricultural soil’ (5 samples), ‘grassland soil’ (4 samples), and ‘farm soil’ (4 samples).

**Table 2.**
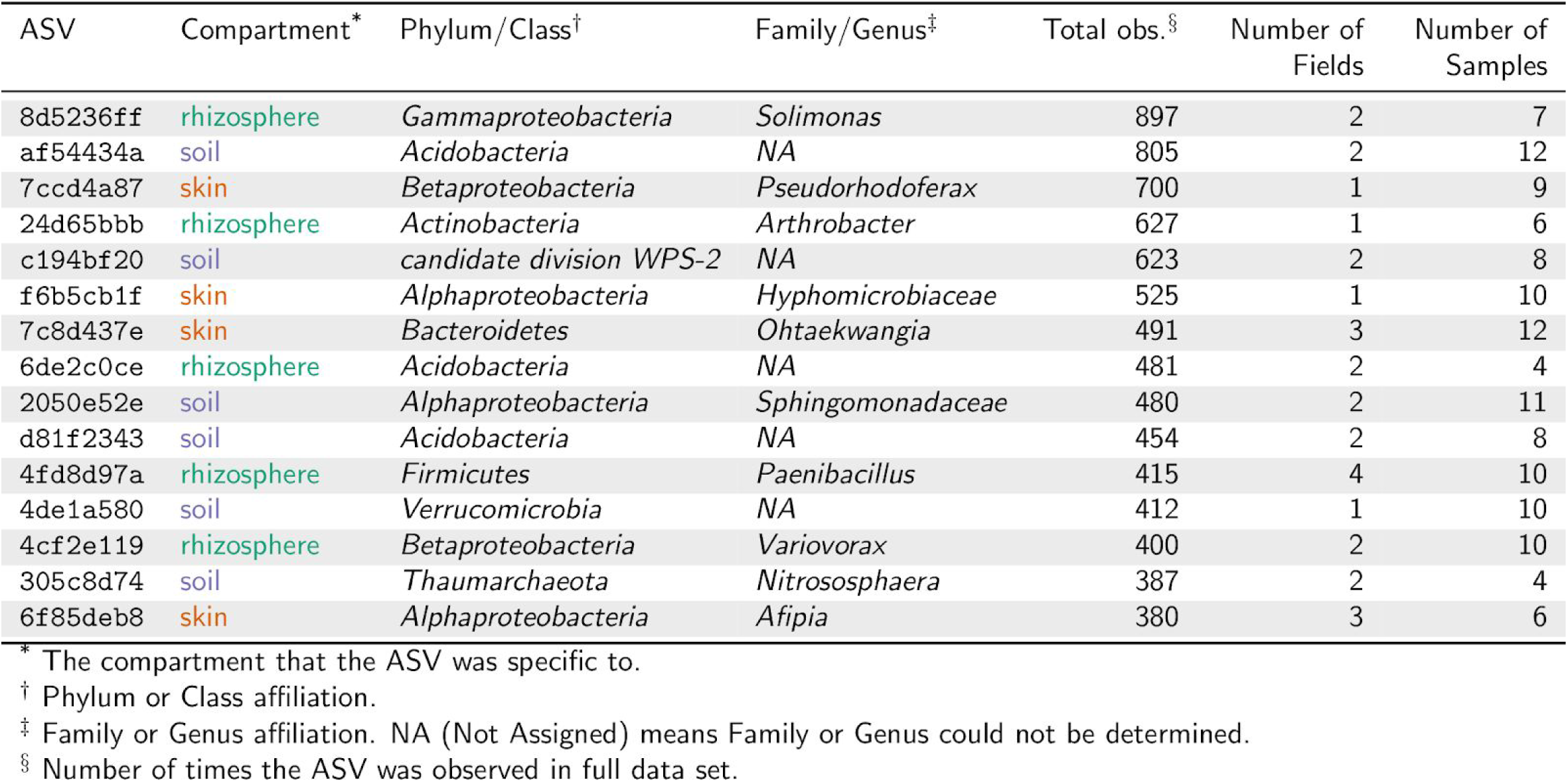
Fifteen topmost compartment-specific ASVs by total sequence count.

| ASV | Compartment* | Phylum/Class† | Family/Genus‡ | Total obs.§ | Number of Fields | Number of Samples |
| --- | --- | --- | --- | --- | --- | --- |
| 8d5236ff | rhizosphere | *Gammaproteobacteria* | *Solimonas* | 897 | 2 | 7 |
| af54434a | soil | *Acidobacteria* | NA | 805 | 2 | 12 |
| 7ccd4a87 | skin | *Betaproteobacteria* | *Pseudorhodoferrax* | 700 | 1 | 9 |
| 24d65bbb | rhizosphere | *Actinobacteria* | *Arthrobacter* | 627 | 1 | 6 |
| c194bf20 | soil | *candidate division WPS-2* | NA | 623 | 2 | 8 |
| f6b5cb1f | skin | *Alphaproteobacteria* | *Hyphomicrobiaceae* | 525 | 1 | 10 |
| 7c8d437e | skin | *Bacteroidetes* | *Ohtaekwangia* | 491 | 3 | 12 |
| 6de2c0ce | rhizosphere | *Acidobacteria* | NA | 481 | 2 | 4 |
| 2050e52e | soil | *Alphaproteobacteria* | *Sphingomonadaceae* | 480 | 2 | 11 |
| d81f2343 | soil | *Acidobacteria* | NA | 454 | 2 | 8 |
| 4fd8d97a | rhizosphere | *Firmicutes* | *Paenibacillus* | 415 | 4 | 10 |
| 4de1a580 | soil | *Verrucomicrobia* | NA | 412 | 1 | 10 |
| 4cf2e119 | rhizosphere | *Betaproteobacteria* | *Variovorax* | 400 | 2 | 10 |
| 305c8d74 | soil | *Thaumarchaeota* | *Nitrososphaera* | 387 | 2 | 4 |
| 6f85deb8 | skin | *Alphaproteobacteria* | *Afipia* | 380 | 3 | 6 |

\* The compartment that the ASV was specific to.
† Phylum or Class affiliation.
‡ Family or Genus affiliation. NA (Not Assigned) means Family or Genus could not be determined.
§ Number of times the ASV was observed in full data set.

Eight of the fifteen ASVs with highest field distribution entropy belonged to the *Alphaproteobacteria* (Table 3). Three belonged to the *Actinobacteria,* one high entropy ASV belonged to the *Firmicutes*, and the rest were other *Proteobacteria*. Only one of the 15 highest entropy ASVs could not be assigned to a taxon at the family rank. This ASV of unknown family membership matches the SSU rRNA gene from *Nordella oligomobilis* strain N21 originally isolated with an amoebal co-culture procedure (La Scola, Barrassi, and Raoult 2004) with 100% sequence identity. The most numerous ASV in the top fifteen ASVs by field distribution entropy matched six EMP ASVs with 100% sequence identity (this is possible as VSearch (Rognes et al. 2016) does a local alignment and any mismatch in the aforementioned comparisons was near ASV sequence ends). The six matching EMP ASVs could be found across 3,286 of the 3,706 ‘soil’ EMP samples (see above).

**Table 3.** Fifteen highest compartment plus field combination entropy ASVs.

| ASV | Phylum/Class* | Family/Genus† | Total obs.‡ | Number of fields§ | Number of samples¶ |
| --- | --- | --- | --- | --- | --- |
| 94e79afc | *Alphaproteobacteria* | *Bradyrhizobium* | 27,134 | 4 | 179 |
| f17e2585 | *Alphaproteobacteria* | *Mesorhizobium* | 14,760 | 4 | 162 |
| 6918660c | *Alphaproteobacteria* | *Devosia* | 19,518 | 4 | 168 |
| 038e3c77 | *Alphaproteobacteria* | NA | 3,055 | 4 | 110 |
| 60493a55 | *Betaproteobacteria* | *Burkholderia* | 3,927 | 4 | 120 |
| 799c493a | *Alphaproteobacteria* | *Phenylobacterium* | 2,270 | 4 | 96 |
| e35a5d71 | *Betaproteobacteria* | *Ramlibacter* | 6,344 | 4 | 91 |
| 57e7ef5d | *Firmicutes* | *Bacillus* | 33,769 | 4 | 197 |
| 14affccc | *Gammaproteobacteria* | *Dokdonella* | 1,079 | 4 | 39 |
| c04d9f02 | *Alphaproteobacteria* | *Methylobacterium* | 2,547 | 4 | 104 |
| dd22ecc6 | *Alphaproteobacteria* | *Bradyrhizobium* | 10,607 | 4 | 100 |
| b4c349f3 | *Alphaproteobacteria* | *Sphingomonadaceae* | 50,955 | 4 | 189 |
| 4064c178 | *Actinobacteria* | *Gaiella* | 3,577 | 4 | 64 |
| 43b7bfbc | *Actinobacteria* | *Aciditerrimonas* | 1,028 | 4 | 39 |
| e96e94b0 | *Actinobacteria* | *Gaiella* | 5,856 | 4 | 123 |

\* Phylum or Class affiliation.
† Family or Genus affiliation. NA (Not Assigned) means Family or Genus could not be determined.
‡ Number of times the ASV was observed in full data set.
§ Number of field locations where the ASV was observed.
¶ Number of samples where the ASV was observed.

### Sweetpotato storage root skin communities were significantly nested by rhizosphere and surrounding soil

Nestedness is a measure of how many community members in less taxa-rich communities are derived from a richer counterpart (see Methods) (Mário Almeida-Neto et al. 2008). In the crop microbiome setting, you might hypothesize, for example, that endophyte communities are derived from -- or ‘nested’ by -- the rhizosphere. Although as the crop develops we might expect nesting to become less pronounced if all compartments diverge from the origin (assuming a common origin). We found sweetpotato skin microbial communities to be significantly nested by rhizosphere communities and soil communities at all sampling locations (p-value 0.01 permutation test, see Methods, Supplementary Figure 6). We did not observe significant nesting of surrounding soil by rhizosphere communities or vice versa. As expected, sweetpotato skin communities did not significantly nest rhizosphere or soil.

### Crop-adjacent bulk soil exhibited the highest ASV alpha-diversity, followed by the rhizosphere, and storage root skin

The crop-adjacent soil was the most ASV-diverse on average relative to the rhizosphere based on the Shannon diversity index (Supplementary Figure 7, p-value □ 0, Tukey’s ‘Honest Significant Difference’ method) and the rhizosphere was richer than skin (p-value □ 0, Tukey’s ‘Honest Significant Difference’ method).

### Beta-diversity decreased with proximity to the sweetpotato

We observed that microbiome composition heterogeneity (beta-diversity) of samples increased with distance from the sweetpotato surface (Figure 5). When using Weighted Unifrac (Lozupone et al. 2011), a phylogeny-aware metric, as the sample distance measure we found beta-diversity to increase from sweetpotato skin to rhizosphere (p-value 6.1e-05, Welch’s two-sample t-test) and from rhizosphere to surrounding soil (p-value 1.9e-15, Welch’s two-sample t-test). This suggests that the sweetpotato skin constrains phylogenetic diversity more than the rhizosphere or soil producing more homogenous microbiome composition among samples.

**Figure 5.**
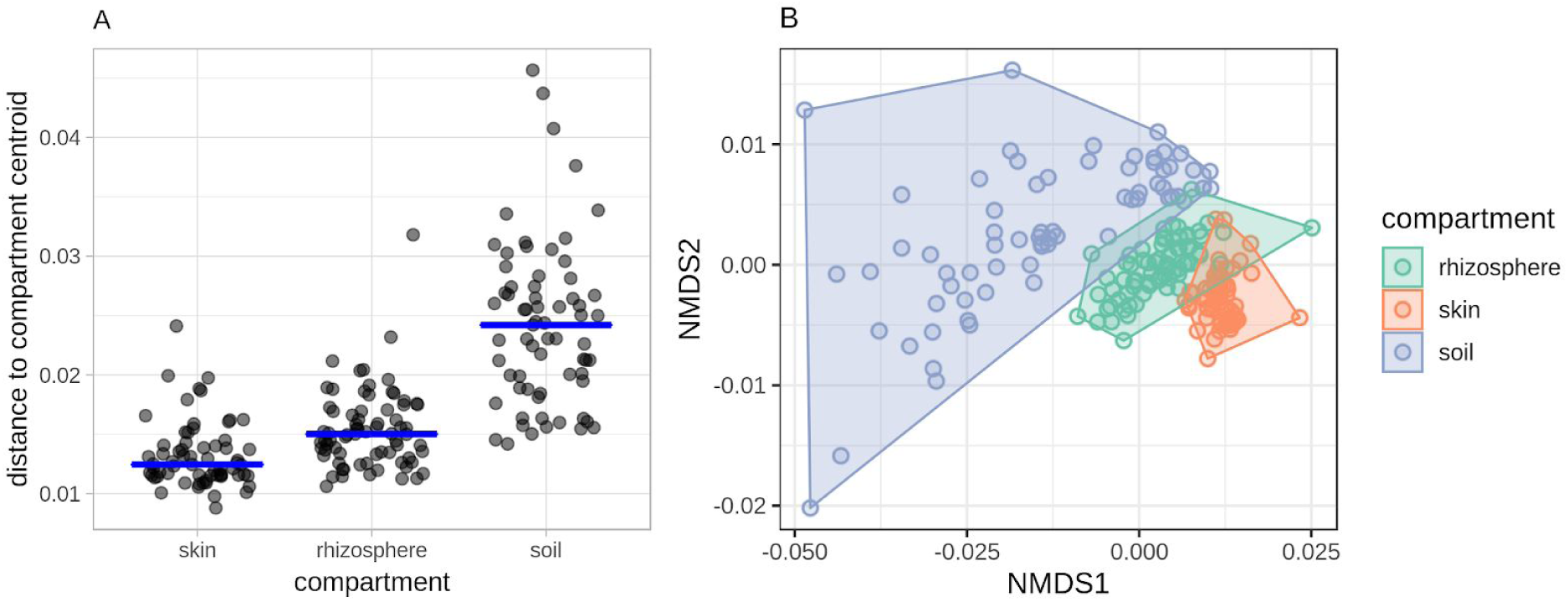
Beta-diversity of Weighted Unifrac distances within compartments. **Panel A** shows the distance to centroid beta-dispersal (Anderson, Ellingsen, and McArdle 2006) values for samples compared with the Weighted Unifrac distance metric (Lozupone et al. 2011) when samples were grouped by compartment. Blue lines indicate the median value. **Panel B** shows the NMDS ordination of Weighted Unifrac distances where points represent samples and color represents compartment. Convex hulls encompass the compartment groups in NMDS-space.

## Discussion

### Sweetpotato rhizosphere and endophytic compartment community assembly

One model for the community assembly of the rhizosphere and plant endophytic compartment hypothesizes assembly happens in two steps (Bulgarelli et al. 2013). In the first step, the soil microbiome community structure in the vicinity of roots shifts in response to plant exudates and other changes in the physiochemical attributes of the soil driven by the plant. In the second step, host (i.e. plant) factors such as the host’s innate immune system select endophytes from the available pool of microorganisms first recruited to the rhizosphere. Four results support the two-step mode for sweetpotato microbiome community assembly: 1) the rhizosphere and endophytic compartments of sweetpotatoes differed significantly from each other and surrounding soil (Figure 2), 2) alpha-diversity decreased significantly from soil to rhizosphere and from rhizosphere to sweetpotato skin (Supplementary Figure 7), 3) rhizosphere microbial community membership significantly nested sweetpotato skin membership (Supplementary Figure 6), and 4) sweetpotato skin possessed the most homogeneous communities in phylogenetic composition followed by the rhizosphere (Figure 5).

The two-step model implies that surrounding soil, rhizosphere, and the endophytic compartment present progressively stronger and different selective pressures to microorganisms, and while all three environments when traced to their origins ultimately start from the soil at planting, different selective pressures favor different microorganisms producing distinct microbial communities. Sweetpotato skin, the rhizosphere, and surrounding soil possessed distinct microbial communities (Figure 2) consistent with the two-step model. Distinct endophyte, rhizosphere, and surrounding soil microbial communities have been observed previously in maize (Peiffer and Ley 2013; Emmett et al. 2017), rice (Edwards et al. 2015), common annual grass (Shi et al. 2015), and *Arabidopsis* (Lundberg et al. 2012), for example.

The two-step model also suggests the endophytic compartment draws from a smaller pool of microorganisms than the rhizosphere and is more selective. Thus, alpha-diversity of the endophytic compartment would be lower than the rhizosphere and surrounding soil under the two-step model, and alpha-diversity of ASVs in this study decreased with proximity to the sweetpotato (Supplementary Figure 7). Similarly to the sweetpotato, Shannon diversity decreased in the endophytic compartment of *Arabidopsis* as compared to surrounding soil (Lundberg et al. 2012), the root communities of three *Arabidopsis* species and *Cardamine hirsuta* possessed generally lower taxa richness than surrounding soil and the rhizosphere (Schlaeppi et al. 2014), and the rhizoplane and endosphere of rice harbored less taxa richness as compared to the rhizosphere and surrounding soil (Edwards et al. 2015).

We did not find other phytobiome assembly studies that quantified nestedness (Mário Almeida-Neto et al. 2008; Thompson et al. 2017) or beta-diversity -- defined as heterogeneity in microbiome composition among samples (Anderson, Ellingsen, and McArdle 2006) --, but it follows from the two-step model of rhizosphere and endophytic compartment community assembly (Bulgarelli et al. 2013) that the endophytic compartment would be nested within the rhizosphere and that beta-diversity would be lowest in the endophytic compartment when compared to the rhizosphere and surrounding soil. That is, under the two-step model, the endophytic compartment is derived from or ‘nested’ by the rhizosphere, and the endophytic compartment is the most selective, followed by the rhizosphere which would produce less heterogeneity in microbiome composition (i.e. beta-diversity) between samples. Sweetpotato skin microbial community membership was significantly nested by rhizosphere at every field location (Supplementary Figure 6). We did not observe statistically significant nesting of the rhizosphere within the soil but we note that at the time of sampling the microbial communities of both surrounding soil and the rhizosphere would have diverged from a common origin and nesting signal would deteriorate over time as the rhizosphere microbial community transitions (Shi et al. 2015).

We observed a decreasing gradient in compartment beta-diversity from the surrounding soil to the rhizosphere to the sweetpotato skin (Figure 5) which suggests sweetpotato skin, of the three environments studied, is the most selective. Disguising oneself from the plant’s immune response or intercepting immune signals (Bulgarelli et al. 2013) may be examples of traits that facilitate microbial endophytic compartment colonization and survival. For example, plant-associated bacterial genomes often contain plant-resembling plant-associated and rhizosphere-associated domains (PREPARADOs) that are hypothesized to disrupt immune signaling in the plant (Levy et al. 2018).

We also observed a number of ASVs that were compartment-specific. Compartment-specific ASVs may mark a profound shift in the abundance rank for the given ASV in one or a combination of compartments from the original community. Compartment-specific ASVs were observed for each compartment. Some were found at only one field location and some spanned multiple locations (Figure 4, Table 2). It is interesting to find putative endophytes not also observed in the adjacent soil or rhizosphere which suggests that microorganisms could be introduced and maintained in the sweetpotato skin in the absence of a persistent reservoir or fitness in the surrounding soil. We cannot say with certainty that sweetpotato skin-specific ASVs are not present the rhizosphere or surrounding soil, but at a minimum they are likely highly enriched in the skin. For example, given how we filtered out ASVs not observed at least ten times and sequenced skin samples to a similar depth as surrounding soil, assuming the next ASV observation in the surrounding soil would be a sweetpotato-skin specific ASV, the least enriched sweetpotato-skin specific ASV would be enriched approximately ten-fold in the skin versus surrounding soil (or the rhizosphere).

Sweetpotato skin-specific ASVs may alternatively indicate vertical transmission of endophytes. Evidence of endophyte vertical transmission has been observed in maize (Johnston-Monje et al. 2014) and other spermatophytes (Shahzad et al. 2018), however, sweetpotatoes are planted as slips for which there is little research regarding routes of vertical transmission. Further research of when and how the sweetpotato acquires endophytes will help to design an approach to engineering the sweetpotato microbiome. The rice endophytic compartment, for context, was shown to quickly acquire microorganisms from surrounding soil and appeared similar in composition to the steady-state endophytic compartment community by thirteen days after planting (Edwards et al. 2015). Rapid endophyte acquisition might enable the introduction of beneficial microorganisms at planting.

### Ecology of taxonomic groups that made up compartment-separating phylogenetic balances

Three phylogenetic balances (Silverman et al. 2017) separated samples by compartment at both farms (balances ‘n5’, ‘n247’, and ‘n138’, Figure 3, Supplementary Figure 2). These three balances represented: **1)** the split between *Acidobacteria* and a super clade of many phyla including *Actinobacteria*, *Firmicutes*, *Gemmatimonadetes,* and *Proteobacteria*, (balance ‘n5’); **2)** the split between *Planctomycetaceae* and a super clade including *Actinobacteria*, *Firmicutes* and *Gemmatimonadetes* (balance ‘n247)’; and **3)** the split between *Alphaproteobacteria* and a narrow clade consisting of undescribed taxa at the phylum level (balance ‘n138’). In fact, the phylogenetic ILR values of these three balances alone encoded enough information to substantively separate sweetpotato skin, rhizosphere, and surrounding soil at both Burch Farm and Sharp Farm (Supplementary Figure 8).

The taxonomic groups captured by the three balances highlighted above have been shown previously to shift in relative abundance across plant compartments; however, a storage root crop like sweetpotato shares little physical resemblance to the root architecture of switchgrass, for instance, and such comparisons should be interpreted with caution. In general, crop-specific meta-analyses may be preferred to control for crop differences. Additionally, there are a number of ways to dissect phytobiomes into their respective compartments and inconsistent methods can produce artifactual patterns of microbiome membership, which could then be falsely attributed to natural phenomena.

Phylogenetic balances ‘n5’, ‘n247’, and ‘n138’ (noted above; Figure 3, Supplementary Figure 2, Supplementary Figure 8) each have one clade (of the two sister clades that makes up each balance) with a strict consensus taxonomic affiliation: *Acidobacteria*, *Planctomycetaceae*, and *Alphaproteobacteria*, respectively. These taxonomic groups have been observed to shift in relative abundance with proximity to roots and/or are widespread, but physiologically undescribed in soil. We summarize relevant features of the *Acidobacteria*, *Planctomycetaceae*, and *Alphaproteobacteria* below:

#### Acidobacteria

*Acidobacteria* have been implicated in the microbial community structure shift between the plant rhizosphere/endosphere and surrounding soil of an annual grass, rice, and *Arabidopsis* (Shi et al. 2015; Edwards et al. 2015; Lundberg et al. 2012). Generally, *Acidobacteria* decrease in relative abundance with proximity to the root although given the nature of compositional data it cannot be determined from SSU rRNA amplicon sequence data how *Acidobacteria* change in *absolute* abundance. It is clear from our results as well as a number of other studies that *Acidobacteria* shift in relative abundance in the rhizosphere when compared to surrounding soil across a wide variety of plants. Soil *Acidobacteria* are considered oligotrophic and/or K-strategists (Fierer, Bradford, and Jackson 2007; Kielak et al. 2016), abundant and widespread (Thompson et al. 2017; Janssen 2006), and difficult to culture with traditional techniques (Kielak et al. 2016).

#### Planctomycetaceae

*Acidobacteria* and *Alphaproteobacteria* are commonly abundant in soil whereas *Planctomycetes* are common but not always abundant (Janssen 2006; Buckley et al. 2006). The family *Planctomycetaceae* are known for unique bacterial intracellular compartmentalization, for members known to carry out ‘anammox’ (the anaerobic oxidation of ammonium), and some *Planctomycetes* are able to metabolize C_1_ compounds (Fuerst and Sagulenko 2011). The ‘n247’ balance value is greater in sweetpotato skin samples than rhizosphere and the numerator clade possesses only *Planctomycetaceae* ASVs which indicates relative enrichment of *Planctomycetaceae* in the endophytic compartment; however, the physiological underpinnings for that phenomenon are unclear given the paucity of known soil *Planctomycetaceae* physiologies. *Planctomycetaceae* were not among isolates from sweetpotato endophyte compartment samples in a previous study (Marques et al. 2015) but this could be attributed to *Planctomycetaceae* resistance to cultivation and/or isolation in the lab. Although the culture-independent method DGGE (denaturing gel gradient electrophoresis) was also used to characterize sweetpotato endophytes, the sequence of only one DGGE band representing one SSU rRNA gene sequence from the community was excised and sequenced (Marques et al. 2015)leaving the specific identity of other endophyte members unresolved.

#### Alphaproteobacteria

*Alphaproteobacteria* are a broad, physiologically diverse class. The most abundant order by ASV affiliation in the *Alphaproteobacteria* clade that makes up one sister clade of balance ‘n138’ is the *Rhizobiales*. The *Rhizobiales* order possesses a number of plant-associated taxa including methylotrophs (Irvine et al. 2012), plant symbionts (Gourion et al. 2015), and plant pathogens (Smith and Townsend 1907). The ‘n138’ balance phylogenetic ILR value was lowest generally in the rhizosphere relative to the sweetpotato skin and especially the surrounding soil (Figure 3, Supplementary Figure 2). *Alphaproteobacteria* abundances make up the denominator of balance ‘n138’ suggesting that *Alphaproteobacteria* are enriched in relative abundance in the rhizosphere and to a lesser extent in the endophytic compartment. The *Alphaproteobacteria* clade of balance ‘n138’ also possesses a number of *Sphingomonadales* ASVs. Both *Sphingomonadales* and *Rhizobiales* have been shown to increase in relative abundance when land is used for agriculture (Souza et al. 2016) and *Sphingomonadales* was among a number of groups shown to previously increase in relative abundance in the sweetpotato rhizosphere (Marques et al. 2014).

### Compartment and field location specific ASVs

Sweetpotato compartments and field locations contained a substantive amount of ASVs specific to the field, the compartment, or both. For instance, 14,418 ASVs were observed at least 10 times and 9,435 of those ASVs were only observed at one field location. Some of these compartment and/or field location-specific ASVs matched closely (at least 99% shared sequence identity) ASVs in the Earth Microbiome Project (EMP) soil data, suggesting a more widespread distribution. However, there were also many compartment and/or field location-specific ASVs that did not match closely any ASV in the EMP soil samples. The effect that field location-specific ASV content has on agroecosystem productivity is unclear. Furthermore, compartment- and/or field location-specific ASVs were generally low in abundance and therefore may be present elsewhere but undetectable at the sequencing depth of this study. Nonetheless, low abundance, possibly compartment- and/or field location-specific ASVs make up a large fraction of the ASV diversity recovered in this study and a smaller but still non-negligible fraction of total ASV observations.

Complicating matters, compartment and/or field location-specific ASVs often did not belong to well-studied phylogenetic groups. Low-abundance members of SSU rRNA gene sequence datasets have been hypothesized to be abundant members of other environments or populations that may become numerically dominant upon environmental change (Sogin et al. 2006). We cannot confirm or eliminate either hypothesis. It may also be that rare, low-abundance ASVs were once numerically dominant and now are derived from ‘relic’ DNA instead of viable microorganisms (Carini et al. 2016). It would be unlikely that the sweetpotato skin specific ASVs came from relic DNA considering the seasonal lifecycle of the sweetpotato. While rare, low-abundance, compartment/field location-specific ASVs are enigmatic, the presence of many sweetpotato skin-specific ASVs suggests there is a wealth of microbial diversity from which to search for plant-beneficial endophytes and promotes a sampling strategy that emphasizes collecting from geographically diverse sites.

## Methods

### Field sampling

We collected samples on September 8th, 2016 from four fields in central North Carolina at two farms, Burch Farm and Sharp Farm. At Burch Farm, we collected samples from three fields; two of which were under a conventional land management regime and one was organically managed. We note there was considerable variation in management practices within the “conventional” and “organic” categories for each field as well. The first conventionally managed field ‘conventional 1’ (35.113054, -78.240148) had ‘Covington’ cultivar sweetpotatoes, the second conventionally managed field ‘conventional 2’ (35.11959, -78.24434) had ‘Beauregard’ cultivar sweetpotatoes, and the organically managed field ‘organic 1’ had two plots, one plot with ‘Beauregard’ (35.10885, -78.23991) and one plot with ‘Covington’ (35.107972, -78.240444). Sweetpotatoes were approximately three months old and near-harvest. We collected from one conventionally managed ‘Covington’ field at Sharp Farm, ‘conventional 3’ (35.70731, -78.12021). Collection sites were spaced approximately ten feet apart and spread across two crop rows to systematically sample the planted area. We brought samples back to the laboratory (AgBiome, Research Triangle Park, NC, USA) on ice and dissected the storage root samples into skin and rhizosphere subsamples. Samples were kept on ice for transport and at 4 C prior to dissection within a week of sampling. Post-dissection, samples were kept at -20 C until DNA extraction. We did not successfully sequence V4 amplicons from three sweetpotato skin samples (one sample each at of the field locations ‘conventional 1’, ‘conventional 3’, and ‘organic 1’). We also collected one fewer crop-adjacent soil sample than sweetpotato storage root sample at field location ‘conventional 2’ and two fewer at field location ‘organic 1’. In total, we sampled and recovered V4 SSU rRNA gene amplicon sequences from 210 samples with an average of 13.8 (s.d. 1.1) crop-adjacent soil, 13.8 (s.d. 0.83) sweetpotato skin, and 14.4 (s.d. 0.9) rhizosphere samples from each cultivar plot.

At Burch Farm, all fields were tilled prior to planting in early June and cultivated during the growing season. Conventionally managed fields were treated with label rates of Lorsban 4E (chlorpyrifos, Dow AgroSciences, Indianapolis, IN) and bifenthrin insecticides, and Valor™ (flumioxazin, Valent, Walnut Creek, CA) and Dual Magnum (S-metachlor, Syngenta) herbicides once per season and received 6-3-18 and 14-4-14 (N-P-K) fertilizers. Organic fields received no treatments other than lime at pre-planting and fields were maintained under organic conditions for approximately five years. Conventional sweetpotato fields at Sharp Farm were tilled prior to planting, fumigated with Telone II (1,3-dichloropropene, Dow AgroSciences, Indianapolis, IN), and treated with a pre-plant incorporation of Lorsban. On May 25th, 2016 sweetpotatoes were planted and fields were treated with Command® (clomazone, FMC, Philadelphia, PA) and Valor herbicides and Brigade® (bifenthrin, FMC) insecticide. At layby, sweetpotatoes received a treatment of Quadris Top (azoxystrobin + difenoconazole, Syngenta) fungicide, Brigade insecticide, and Dual Magnum herbicide and then were treated in August with Endigo® ZC (lambda-cyhalothrin + thiamethoxam, Syngenta) and Coragen® (chlorantraniliprole, Dupont, Wilmington, DE) insecticides.

### DNA extraction, PCR, and DNA sequencing of V4 amplicons

Rhizosphere samples were collected by washing and collecting soil attached to the sweetpotato storage root samples with 0.2 μm filtered ddH_2_0. Sweetpotato storage root skin from sterilely washed sweetpotato storage roots was removed with a razor, ground with a mortar and pestle, and collected in a slurry with 0.2 μm filtered phosphate buffered saline. DNA was extracted from sweetpotato skin, rhizosphere, and soil samples via the MO Bio DNeasy PowerSoil HTP 96 Kit (product number 12955-4) using the manufacturer’s protocol. SSU rRNA gene V4-region DNA was PCR amplified using ‘modified’, up-to-date, and improved V4 SSU rRNA gene primers (Walters et al. 2016). PCR was done on a Fluidigm Digital PCR instrument (model FC1) with alternating conventional denaturation/annealing/elongation and C_0_t cycles (denaturation/C_0_t/annealing/elongation). Specifically, PCR began with a thermal mix step (50 C 120 sec.); a hot start (90 C 600 sec.); 10 cycles of denature (95 C 15 sec.), annealing (55 C 30 sec.), and elongation (72 C 60 sec.); 2 cycles of denature (95 C 15 sec.), C_0_t (80 C 30 sec.), annealing (55 C 30 sec.), and elongation (72 C 60 sec.); 8 cycles of denature (95 C 15 sec.), annealing (55 C 30 sec.), and elongation (72 C 60 sec.); 2 cycles of denature (95 C 15 sec.), C_0_t (80 C 30 sec.), annealing (55 C 30 sec.), and elongation (72 C 60 sec.); 8 cycles of (95 C 15 sec.), annealing (55 C 30 sec.), and elongation (72 C 60 sec.); 5 cycles of denature (95 C 15 sec.), C_0_t (80 C 30 sec.), annealing (55 C 30 sec.), and elongation (72 C 60 sec.); and a final elongation (72 C 180 sec). All PCR reagents were part of the Roche FastStart High Fidelity PCR System (product number: 3553400001). Amplicons were sequenced on the Illumina MiSeq and prepared for sequencing with the Illumina v3 chemistry (600-cycle, product number: MS-102-3003). Forward primers included a 10 bp index specific to each sample to demultiplex reads by sample post-DNA sequencing. All raw amplicon sequences are available at the European Nucleotide Archive (study accession: PRJEB29367, https://www.ebi.ac.uk/ena/data/view/PRJEB29367)

### Amplicon sequence read quality control (QC) and preprocessing

We removed primer sequences, quality trimmed, corrected errors, and merged paired V4 SSU rRNA gene amplicon reads with the DADA2 (Callahan et al. 2016) pipeline wrapped by QIIME2 (Caporaso et al. 2010). Specifically, the QC pipeline trimmed 25 nt from the beginning forward and reverse reads to excise primer sequences. We also truncated forward and reverse reads at position 230 (forward read) and 180 (reverse read) to discard read regions where quality scores appeared to drop (Supplementary Figure 9). Following trimming, we removed any reads for which the number of expected errors (Edgar and Flyvbjerg 2015) exceeded 1. We selected to train the DADA2 error model with 1,000,000 reads from each run (a separate model was trained for each sequencing run), and following error correction, we merged reads with default DADA2 parameters (minimum overlap of 12 nt, zero mismatches allowed). Chimeras were identified with the ‘consensus’ DADA2 method and discarded. We used version 1.6.0 of the DADA2 R package and version 2017.12 of QIIME2. We began with 8,327,842 raw DNA sequence reads (an average of 34,845 per sample (s.d. 8,106)) and following QC the full data set contained 4,371,412 merged amplicon sequences, (an average of 18,290 per sample (s.d. 5,569)). The QC pipeline output consisted of amplicon sequence variant (ASV) counts (Callahan, McMurdie, and Holmes 2017) in a Phyloseq R object (McMurdie and Holmes 2013). We stitched together all QC pipeline operations with the Nextflow bioinformatics workflow DSL (domain specific language) (Di Tommaso et al. 2017).

Following read merging, quality and primer trimming, and error correction, we discarded reads that best-matched sequences in the SILVA SSU rRNA gene database (SILVA v123) (Quast et al. 2013) that were annotated as “Mitochondria”, “Chloroplast”, or “Eukarya”. We also discarded any samples from further analysis for which we did not recover at least 5,000 amplicon sequences leaving 204 samples. There were 22,090 ASVs in our final dataset and 2,925,851 total amplicon observations for an average of 14,342 per sample.

In addition to experimental samples, we sequenced V4-region SSU rRNA gene from five replicates of a mock community (ZymoBIOMICS; Irvine, CA; Microbial Community Standard: product number D6305). The mock community possessed genomic DNA from eight bacteria and two fungi: *Pseudomonas aeruginosa, Escherichia coli, Salmonella enterica, Lactobacillus fermentum, Enterococcus faecalis, Staphylococcus aureus, Listeria monocytogenes, Bacillus subtilis, Saccharomyces cerevisiae, Cryptococcus neoformans*. We recovered thirteen total ASVs from the mock community DNA. One mock community ASV appeared to be a non-target, non-SSU rRNA gene sequence product matching with 100% sequence identity several genomes of *Lactobacillus Fermentum* and was observed 14 times in only one sample accounting for in 7.2E-6% of the 19,544 total observations from that sample. Another ASV belonged to *Arthrobacter* but was only found in one sample and observed only 3 times in 20,751 total observations from that sample (1.4E-6% of total observations). The remaining ASVs could all be attributed to the expected mock community members (Supplementary Figure 10).

### Phylogenetic Tree

We aligned quality-controlled V4 SSU rRNA gene amplicon sequences against the ‘bacterial’ covariance model provided with the ‘ssu-align’ aligner version 0.1.1 (Nawrocki 2009). Alignment positions that were not accounted for in the covariance model were discarded. We further discarded columns where less than 95% of nucleotides had alignment posterior probability less than 95%. The final alignment included 258 positions.. The phylogenetic tree was constructed using the general time reversible model for nucleotide evolution and the FastTree version 2.1.10 approximately-maximum-likelihood algorithm (Price, Dehal, and Arkin 2010, 2009). We constrained the tree topology to be consistent with phylum affiliation of the SSU rRNA gene amplicon sequences. The tree was rooted at the branch that differentiated *Archaea* from *Bacteria*. Post-rooting, all tips that were more than three orders of magnitude longer than the mean tip distance from the root were assumed anomalous and the corresponding ASVs were discarded from the dataset. Post-filtering, 21,608 ASVs remained.

### Assessing taxonomic affiliation of amplicon sequence reads

We determined the taxonomic affiliation of amplicon sequence reads using the Ribosomal Database Project (RDP) naive Bayesian classifier algorithm (Wang et al. 2007) as implemented in the DADA2 R package (Callahan et al. 2016) with a minimum bootstrap confidence threshold of 50 and trained on version 16 of the RDP SSU rRNA gene sequence database (Cole et al. 2009). Species-level assignments were only given for sequences that perfectly matched only a single reference as proposed recently (Edgar 2017). The reference sequences for determining species-level taxonomic affiliation were provided by the developers of the DADA2 R package (https://benjjneb.github.io/dada2/training.html) and were originally derived from the RDP database version 16.

### Ordination of sample microbial community composition

We measured differences in the microbial composition of each sample with the Bray-Curtis (Bray and Curtis 1957) metric or the Weighted Unifrac Metric (Lozupone et al. 2011). We used Non-metric-multidimensional scaling (NMDS) (Kruskal 1964) for dimensionality reduction and testing the significance of microbial community composition differences across sample groups (Oksanen et al. 2013; Anderson 2001). Beta-diversity is a measure of the compositional heterogeneity between samples and the beta-diversity for a group of samples can be quantified from a sample pairwise distance matrix. A centroid in multivariate space is calculated given the pairwise distances of samples within a category group (e.g. ‘rhizosphere’ samples). Individual samples within the group are compared to the group centroid yielding a distance from the group centroid for each sample (Anderson, Ellingsen, and McArdle 2006). In this study, the distance for a sample from the group centroid in multivariate space was calculated from Weighted Unifrac sample pairwise distance matrices (Oksanen et al. 2013; Anderson, Ellingsen, and McArdle 2006).

### Quantifying relative representation of microbial community composition variance by land management, sweet potato cultivar, and region

To assess the relative representation of microbial community variance by sample characteristics within a single farm, we first transformed our relative ASV abundance data to ‘balances’ (Morton et al. 2017) or specifically isometric log abundance ratios (ILR) of descendant ASVs at nodes of a bifurcating ASV graph. The bifurcating graph was created by clustering sample ASV abundance profiles across samples by the Ward agglomerative technique (Ward 1963). Low abundance ASVs were filtered from the data prior to calculating the ILR transform. Specifically, we removed any ASV that was not present at least twice in at least 15% of samples. After sparsity filtering, there were 363 total ASVs in the data. Our regression model included terms representing crop compartment microbiome comparisons (i.e. sweetpotato storage root ‘skin’ versus surrounding ‘soil’ and ‘rhizosphere’ versus surrounding ‘soil’), the field within the farm where the sample came from, and the sweetpotato cultivar associated with the sample (i.e. ‘Covington’ or ‘Beauregard’). We dummy-encoded the sample compartment variable (e.g. ‘soil’, ‘skin’, or ‘rhizosphere’) using ‘soil’ as the reference category, the field category using the organic field as the reference, and the cultivar category using ‘Covington’ as the reference. We modeled balances with sample variables using multivariate, multiple ordinary least squares regression (using functions of the compositional data analysis Python package ‘gneiss’ (Seabold and Perktold 2010; Morton et al. 2017)). To assess the relative contribution of each sample metadata category to the model’s predictive value, we used a leave-one-out approach where we calculated the difference in the coefficient of multiple determination for models upon leaving out each possible model term separately (Morton et al. 2017).

### Calculating phylogenetic isometric log ratios

Phylogenetic ILR values for balances were made by weighting taxa using the geometric mean times the Euclidean norm which assigns less weight to taxa with more zero and near-zero counts. Before calculating phylogenetic ILR values we removed low abundance ASVs. That is, we removed any ASV not observed at least twice in 13.5% percent of all samples across Burch Farm and Sharp Farm leaving 418 ASVs. Filtering criteria was changed from the *single* farm analysis (see above) -- i.e. from ASVs observed at least twice in at least 15% of samples to ASVs observed at least twice in at least 13.5% of samples -- to incorporate more ASVs for the phylogenetic ILR-based analysis which included both Burch and Sharp Farm as opposed to just Burch. We weighed phylogenetic balances using the square root of summed branch lengths. Our taxa and weighting scheme mimicked exactly the scheme wherein the phylogenetic ILR transform was defined (Silverman et al. 2017). All phylogenetic balances were calculated with the R package written by the authors for the phylogenetic ILR transform ‘philR’ (Silverman et al. 2017).

### Identifying phylogenetic balances that separate samples from different crop compartments

We fit a multinomial, penalized, logistic regression model (Friedman, Hastie, and Tibshirani 2009) of compartment categories with phylogenetic ILR values (see Silverman et al. (2017) for a thorough description of the phylogenetic ILR transform and subsequent modeling). The logistic regression penalization term *lambda* was set at 0.045 after inspecting how nonzero balance coefficients appeared to separate samples by compartment. The models included 25, and 13 nonzero regression coefficients for Burch Farm and Sharp Farm, respectively..

### Calculating ASV distribution entropy

ASV distribution entropy was calculated from frequency vectors of ASV observations into sample categories (Thompson et al. 2017). Depending on the analysis, we either grouped samples into compartment categories or combinations of compartment plus field location categories. Entropy was calculated using the natural logarithm. Briefly, entropy in this context measures the heterogeneity of a taxon’s distribution across sample groups. Low entropy means a taxon (ASV or genus) is found in few groups whereas high entropy indicates a taxon is found in many groups evenly

### Comparing ASVs in this study to Earth Microbiome Project soil ASVs

We downloaded the Earth Microbiome Project (EMP) (Thompson et al. 2017) ASV data from ftp://ftp.microbio.me/emp/release1/. We selected the ‘`emp_deblur_90bp.qc_filtered.biom`’ version of the EMP data which includes the unrarefied data used by Thompson *et al*. (2017). Specifically, the full EMP dataset included 23,828 samples and 317,314 ASVs. From the full data, we selected any sample for which the ‘`env_material`’ value was either ‘soil’, ‘bulk soil’, or ‘rhizosphere’.. We filtered out any ASV with a total of zero counts across all samples leaving 160,491 ASVs from a broad geographic distribution of samples. We searched for EMP matches to the ASVs generated in this study using VSearch with a sequence identity threshold of 99% and both ‘maxaccepts’ and ‘maxrejects’ set to zero (Rognes et al. 2016).

### Measuring nestedness between sample compartments

We measured nestedness using the ‘nestedness metric based on overlap and decreasing fill’ (nestedNODF) method as implemented in the R package *Vegan* (Oksanen et al. 2013; Mario Almeida-Neto et al. 2008), however, we only considered the N_rows_ value -- which we refer to as “**N_samples_**” -- which corresponds to just sample nesting as opposed to taxa nesting or the two-dimensional nesting of the full sample by taxon presence/absence matrix. Prior to calculating nestedness, samples were ordered by compartment affiliation as the first ordering priority and the number of taxa observed as the second ordering priority with more rich samples ordered above less rich samples. The sample order allowed us to measure the extent that one compartment nested another. To assess the statistical significance of N_samples_ values, we randomly permuted sample compartment labels 100 times and found how often N_samples_ values from permuted data exceeded that of the measured N_samples_ value with true labels.

## Acknowledgments

We are grateful to Mauricio J. Borgen (AgBiome) for scientific computing support, MOGene LLC (St. Louis, MO) for DNA sequencing services, and Burch Farm and Sharp Farm for generously letting us collect samples. We especially thank the R/Python package developers of the many open-source packages used in this study without which data-intensive biology would not be possible. This work was funded by a grant from the Bill and Melinda Gates Foundation (Investment ID OPP1134622). We thank Jeffery L. Dangl and Isai Salas González for clarifying edits and useful discussion. Also, a number of employees at AgBiome made helpful edits and improvements to the manuscript including Dr. Alexandra B. Crawley, Dr. Matthew B. Biggs, Dr. Dan J. Tomso, and Dr. Eric Ward.

## Supplementary Figures

**Supplementary Figure 1.**
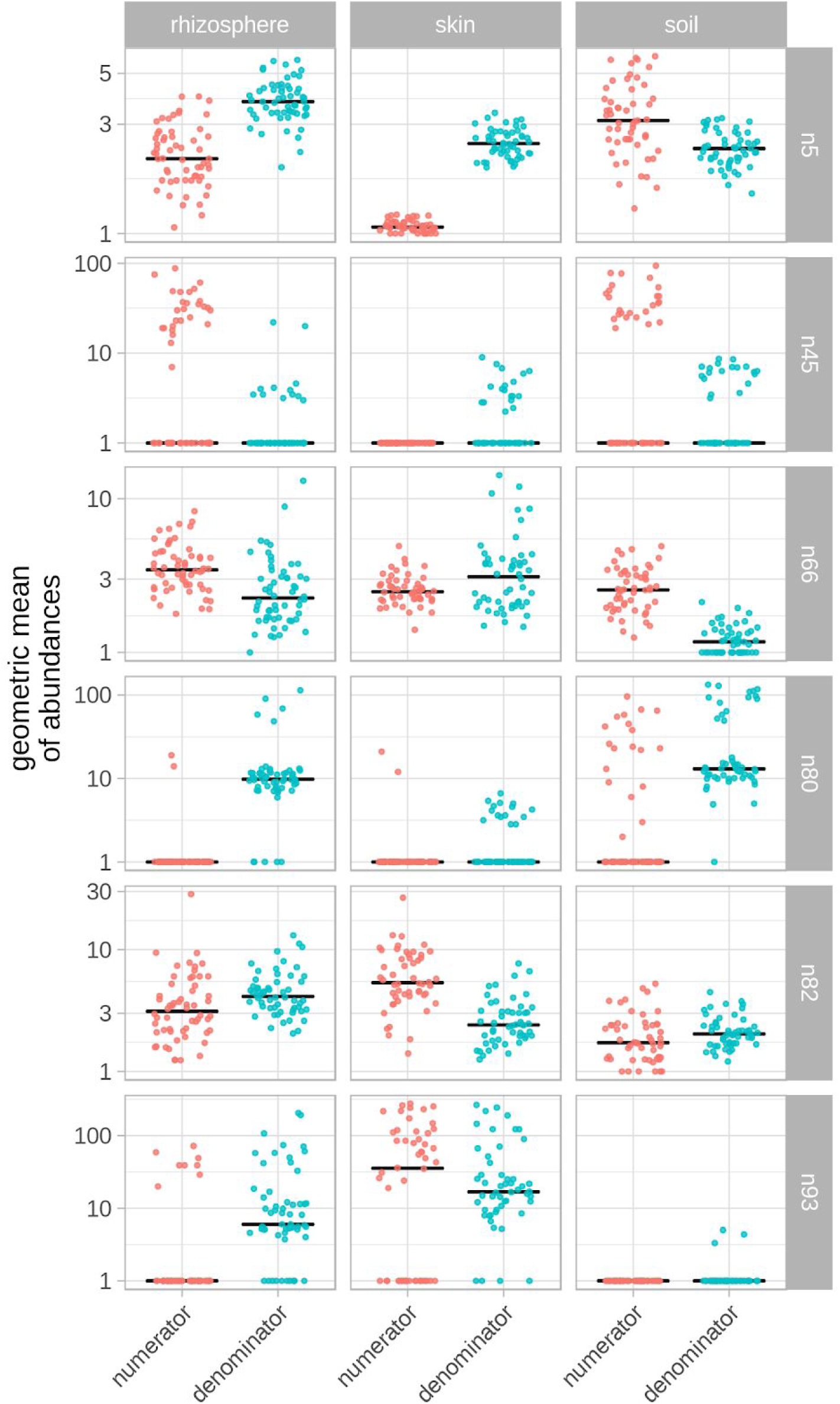
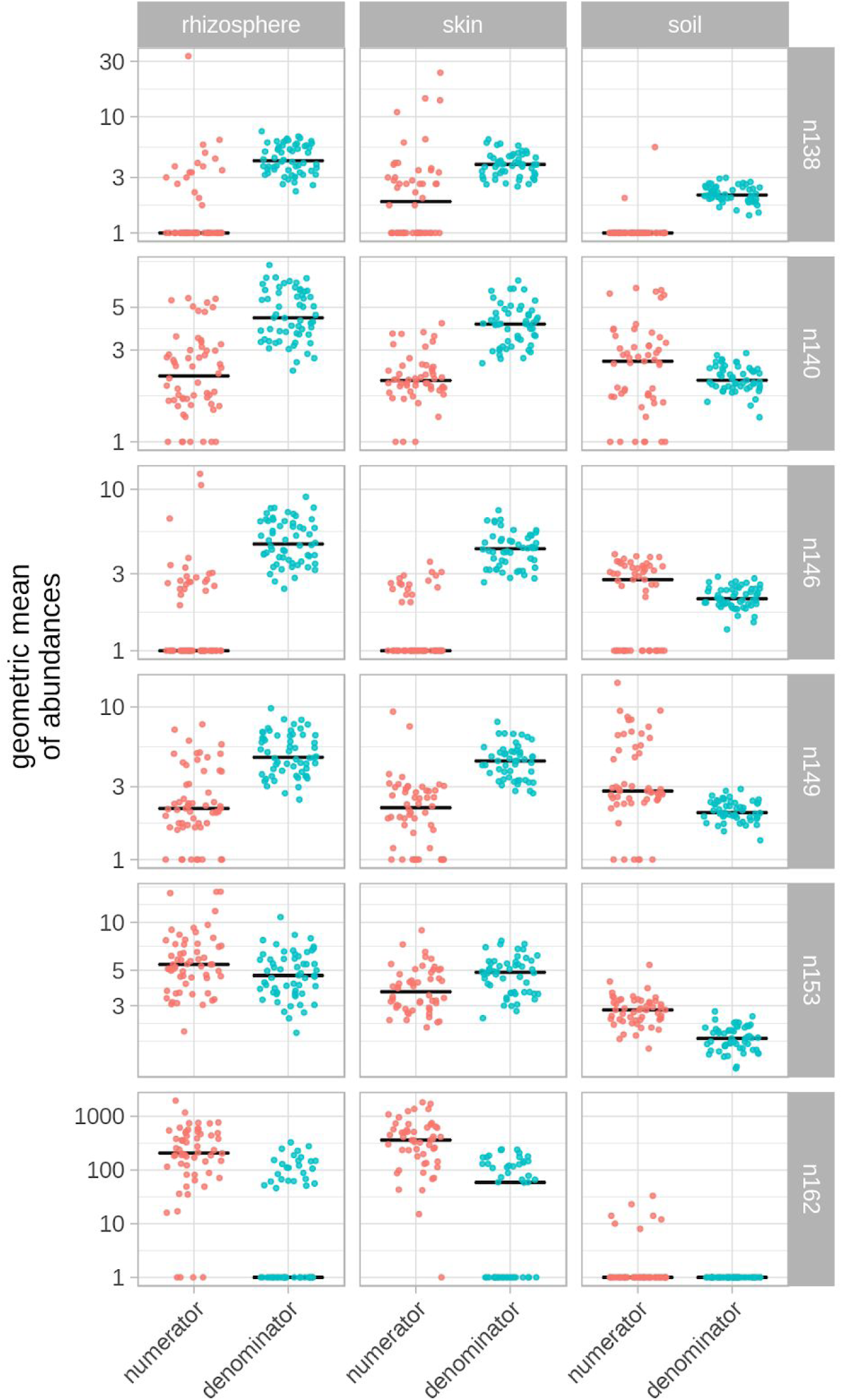
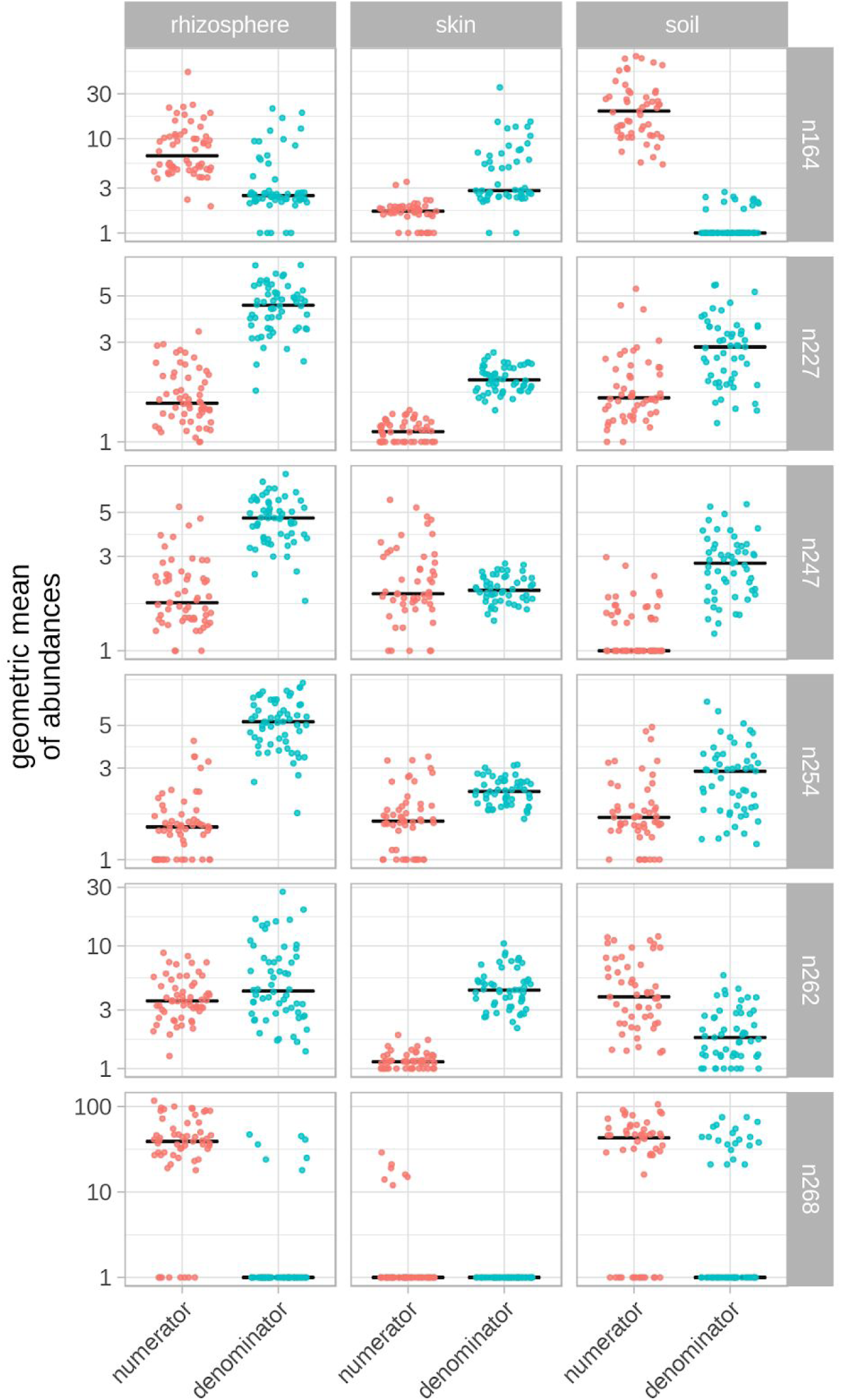
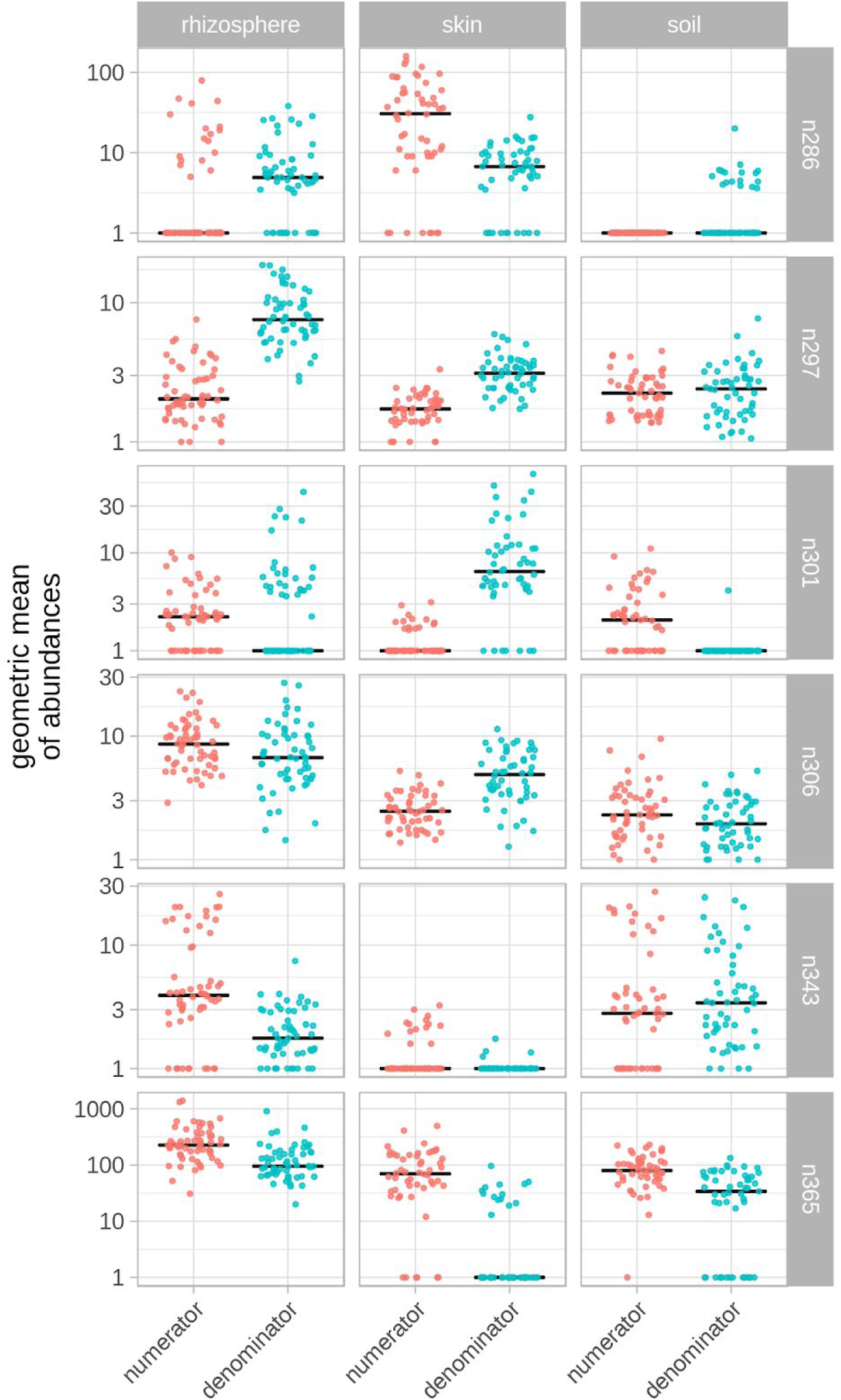
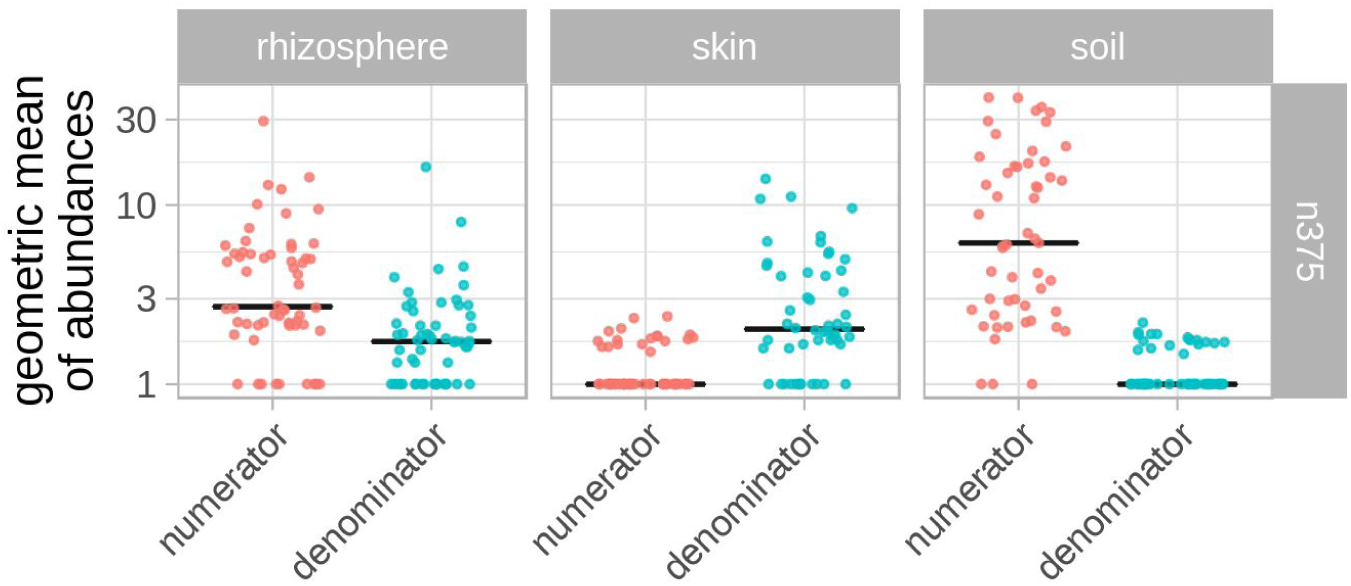
Geometric mean of balance sister clade abundances within a sample (Burch Farm) Each point represents the geometric mean of counts within a sample for all ASV members of either the numerator or denominator clade of the given balance. Every row represents an individual balance labeled on the right. Each cell represents one compartment-balance combination. Compartment labels are shown across the top of the figure. See Figure 3 for the phylogenetic ILR values and positions of each balance. Colors are consistent with the numerator/denominator colors in Figure 3. The black line denotes the median value.

**Supplementary Figure 2.**
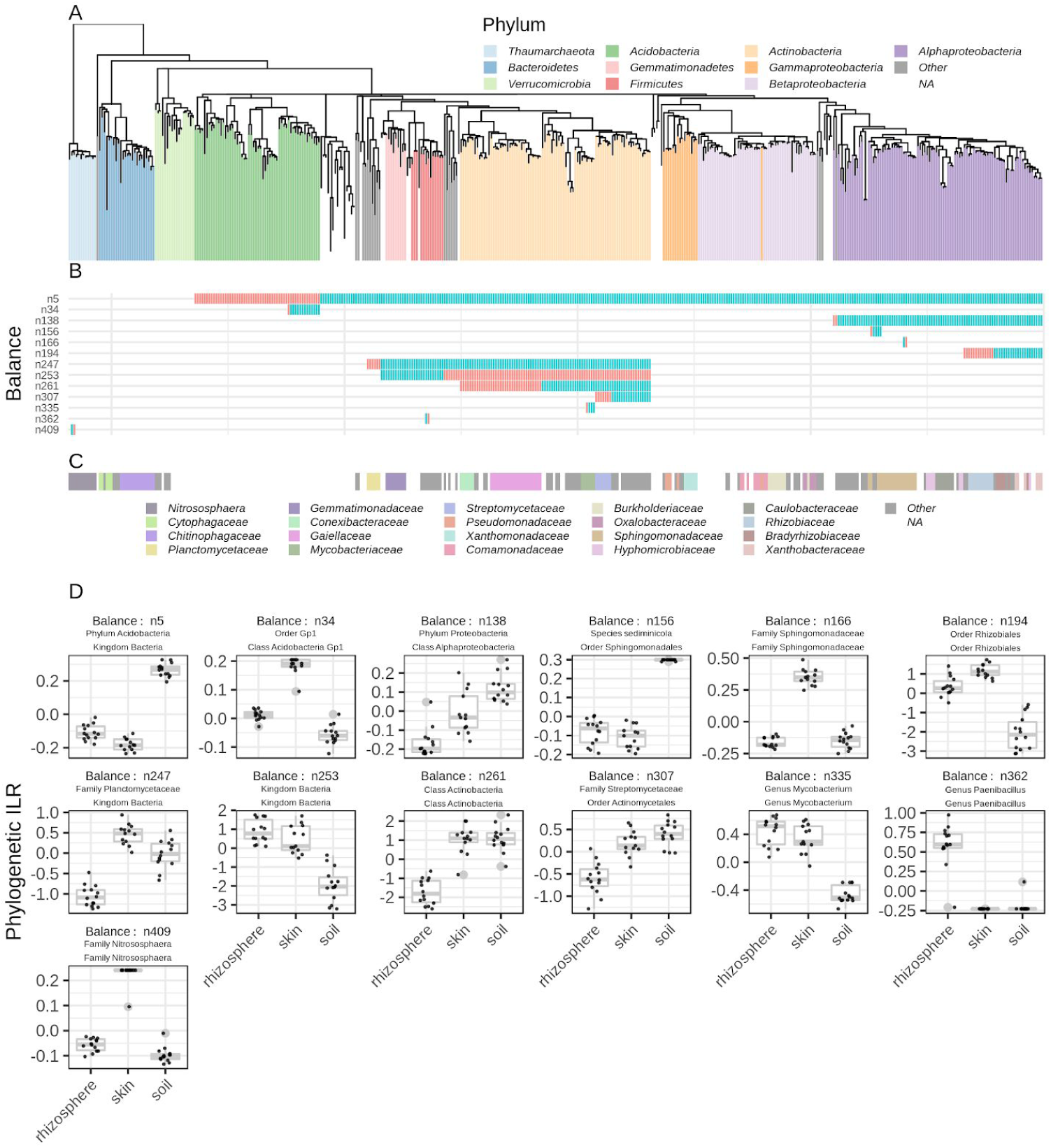
Phylogenetic balances that separated samples by compartments at Sharp Farm. Position and phylogenetic ILR (Silverman et al. 2017) values for balances found to separate samples by compartment at **Sharp Farm**. **Panel A** shows the phylogenetic relationship of ASVs used in the phylogenetic balance analysis (see Methods). **Panel B** shows the positioning in the phylogeny of each balance. Colors in the second panel denote which side (i.e. sister clade) of the balance was used as the denominator (blue) or numerator (red) when calculating the phylogenetic ILR for each balance. **Panel C** shows the position of Family-level taxonomic groups in the tree. **Panel D** shows the balance ILR values grouped by compartment for each balance highlighted in the second panel. Taxa names in the title of each **Panel D** subplot are the highest resolution, 95% consensus taxonomic affiliation for the sister clades that makes up the balance. The consensus taxonomy of the sister clade used as the numerator to calculate the balance phylogenetic ILR is listed first followed by the denominator clade. Boxplot boxes extend from the 25th to the 75th percentile and include a horizontal line indicating the median. Whiskers extend to the largest value that is no greater than 1.5 times the interquartile range from the 25th (bottom of the box) or 75th (top of the box) percentile. Points beyond the whiskers are plotted individually.

**Supplementary Figure 3.**
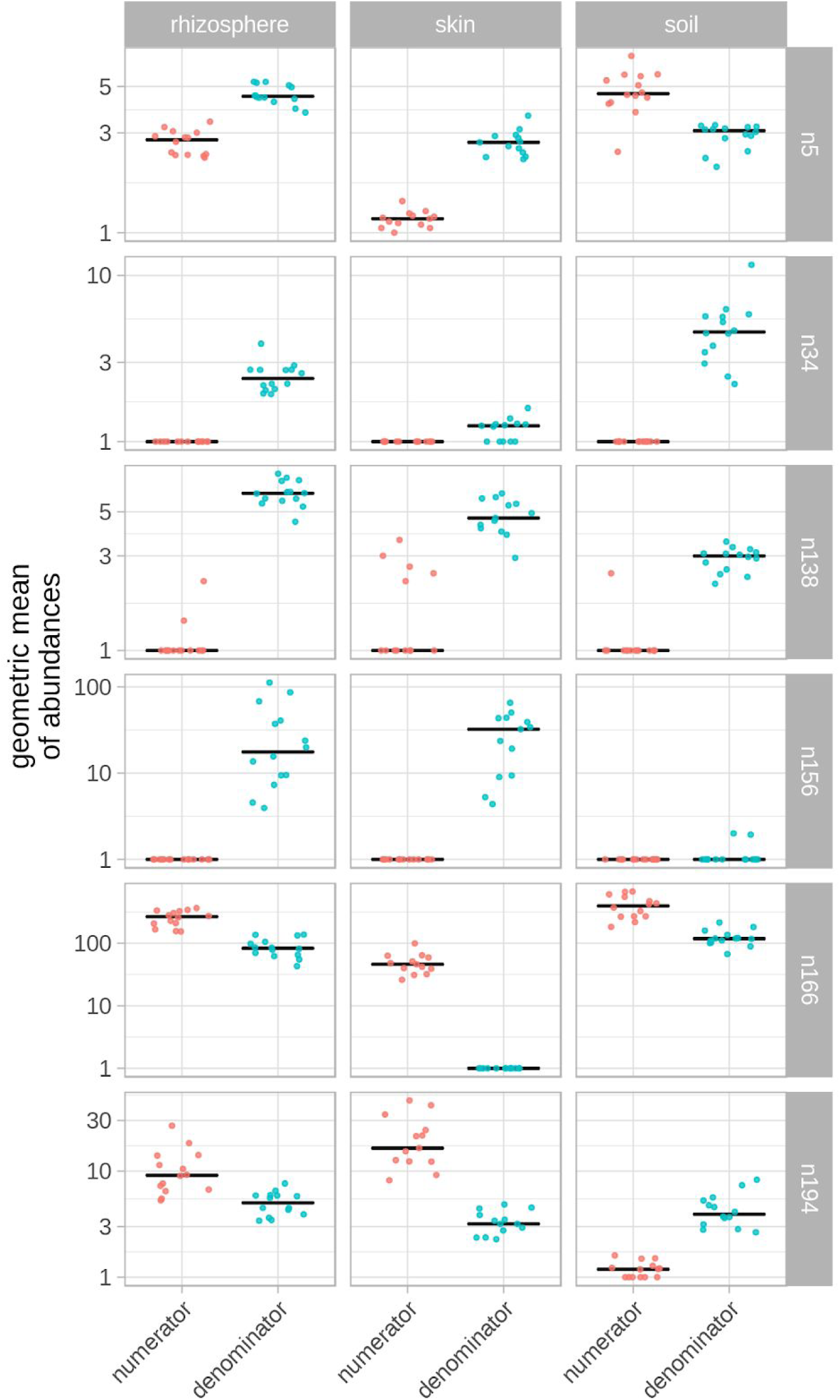
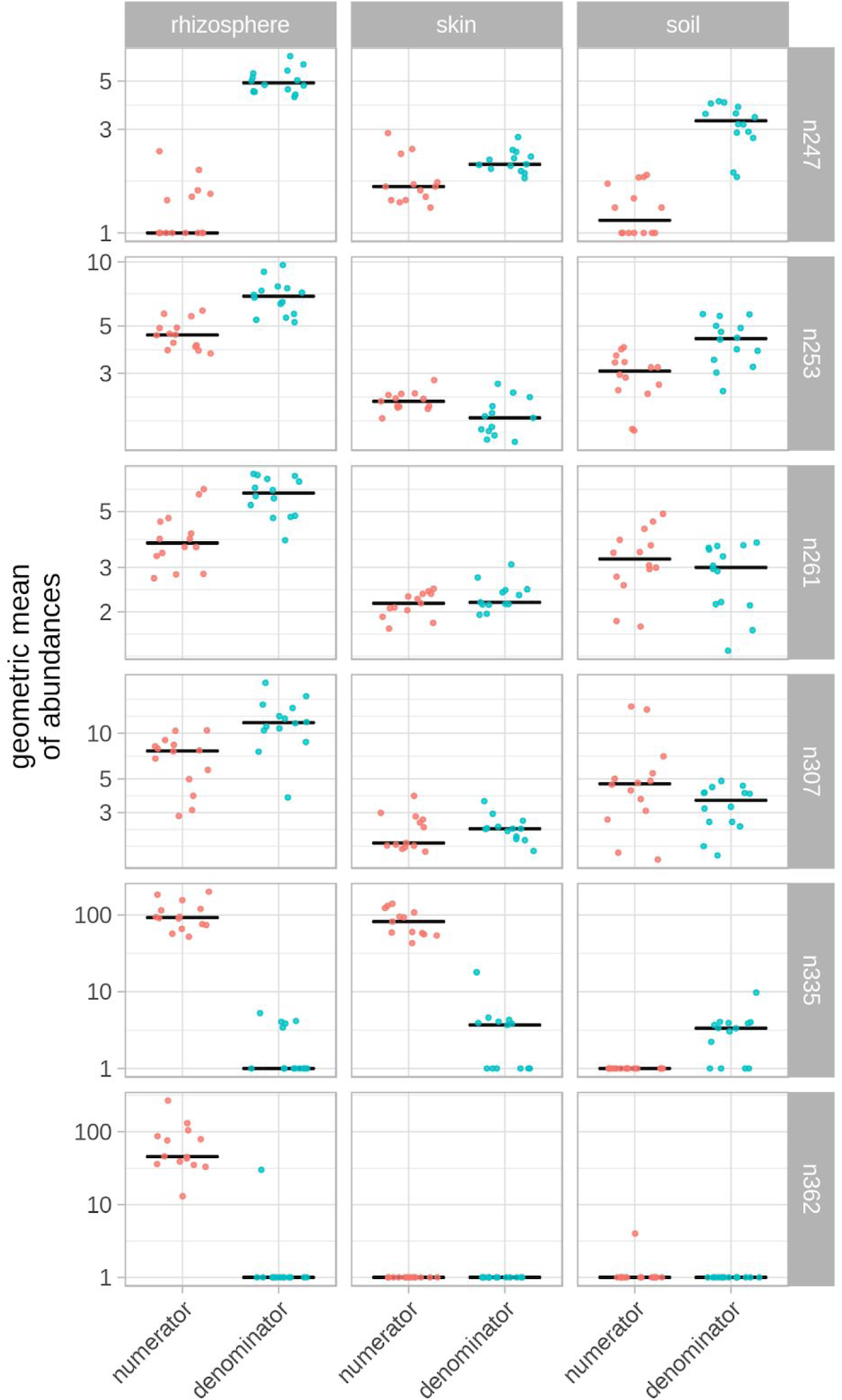
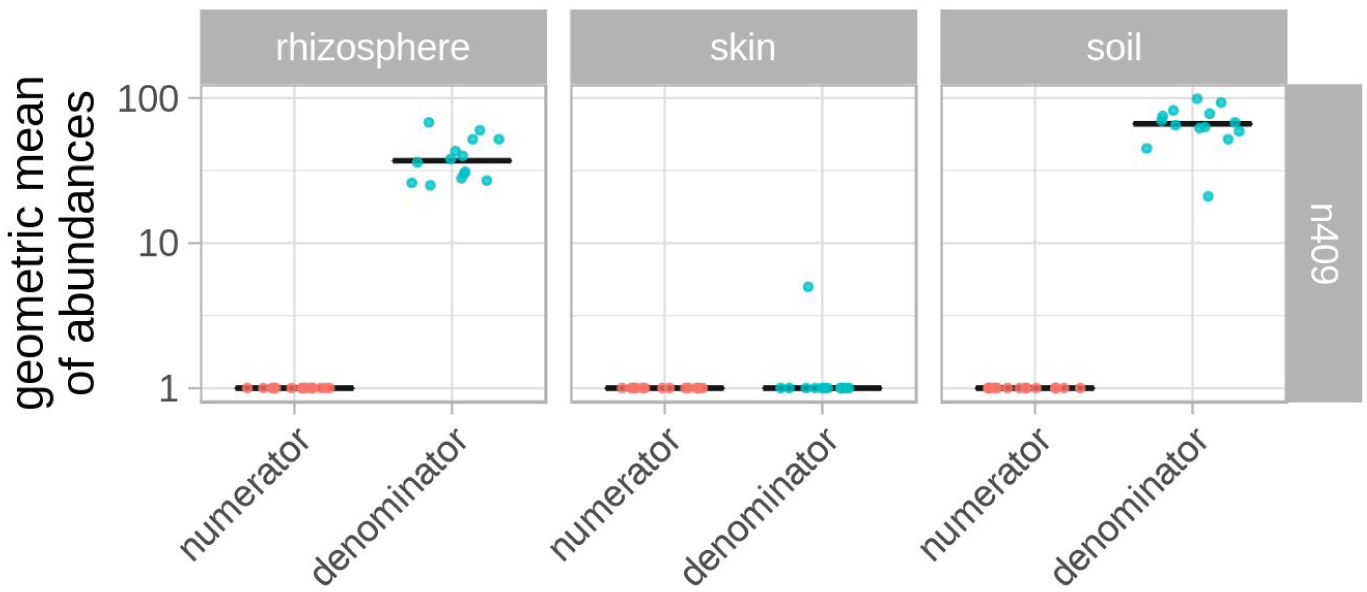
Geometric mean of balance sister clade abundances within a sample (Sharp Farm) Each point represents the geometric mean of counts within a sample of ASV members within either the numerator or denominator clade of the given balance. Every row represents an individual balance labeled on the right. Each cell represents one compartment-balance combination. Compartment labels are shown across the top of the figure. See Supplementary Figure 2 for the phylogenetic ILR values and positions of each balance. Colors are consistent with the numerator/denominator colors in Supplementary Figure 2. The black line denotes the median value.

**Supplementary Figure 4.**
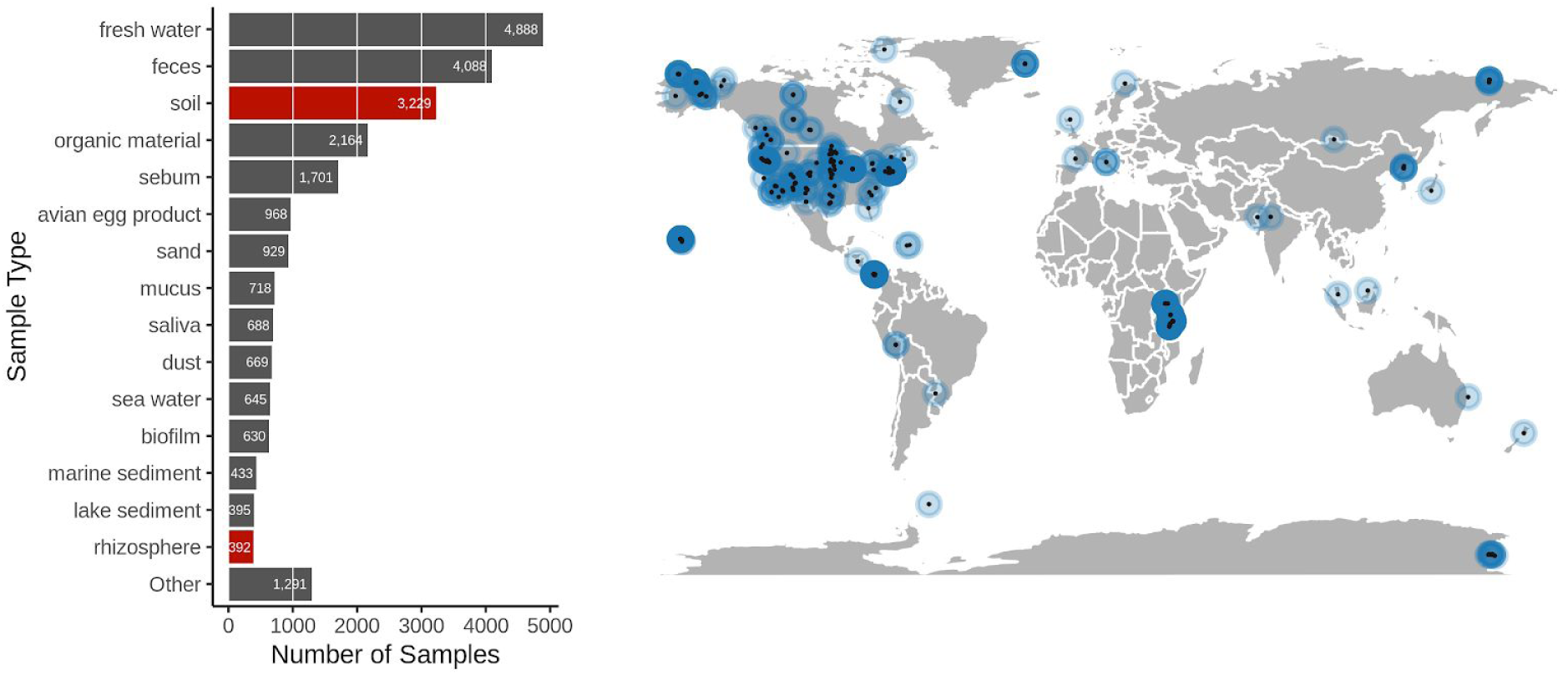
Earth Microbiome Project samples used in this study for context. Locations of Earth Microbiome Project (EMP) samples used for reference in this study (**right panel**). Bar chart of the 15 most numerous ‘`env_material`’ values in the EMP project (**left panel)**. The bars in red indicate which samples were selected for reference for this study. We also included the ‘bulk soil’ ‘`env_material`’ value but it was not in the fifteen most numerous values.

**Supplementary Figure 5.**
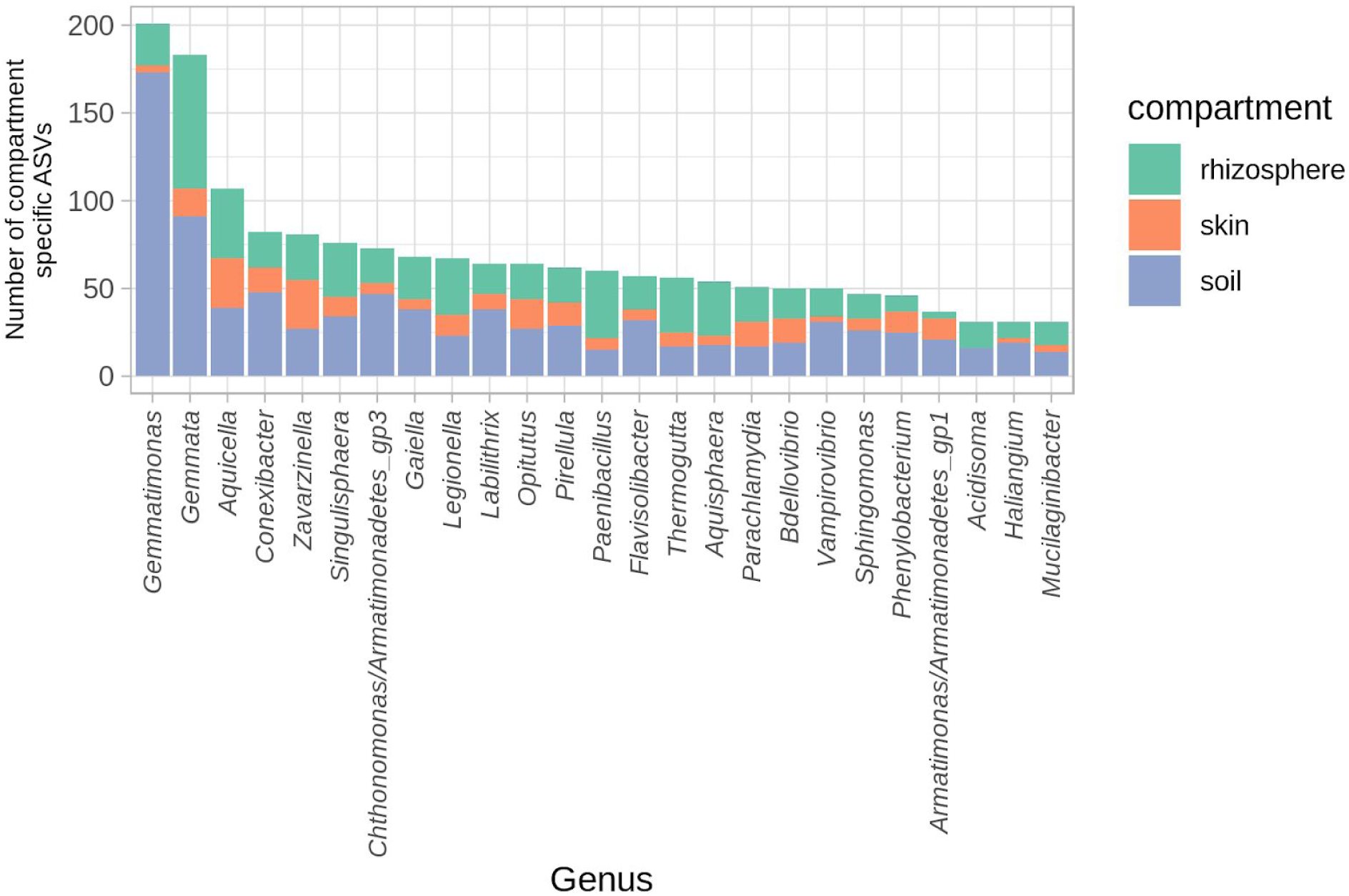
Genus affiliation of compartment-specific ASVs. Genus affiliation of compartment-specific ASVs. Color denotes the compartment where the ASV is specifically found. ASVs without a genus affiliation are not shown.

**Supplementary Figure 6.**
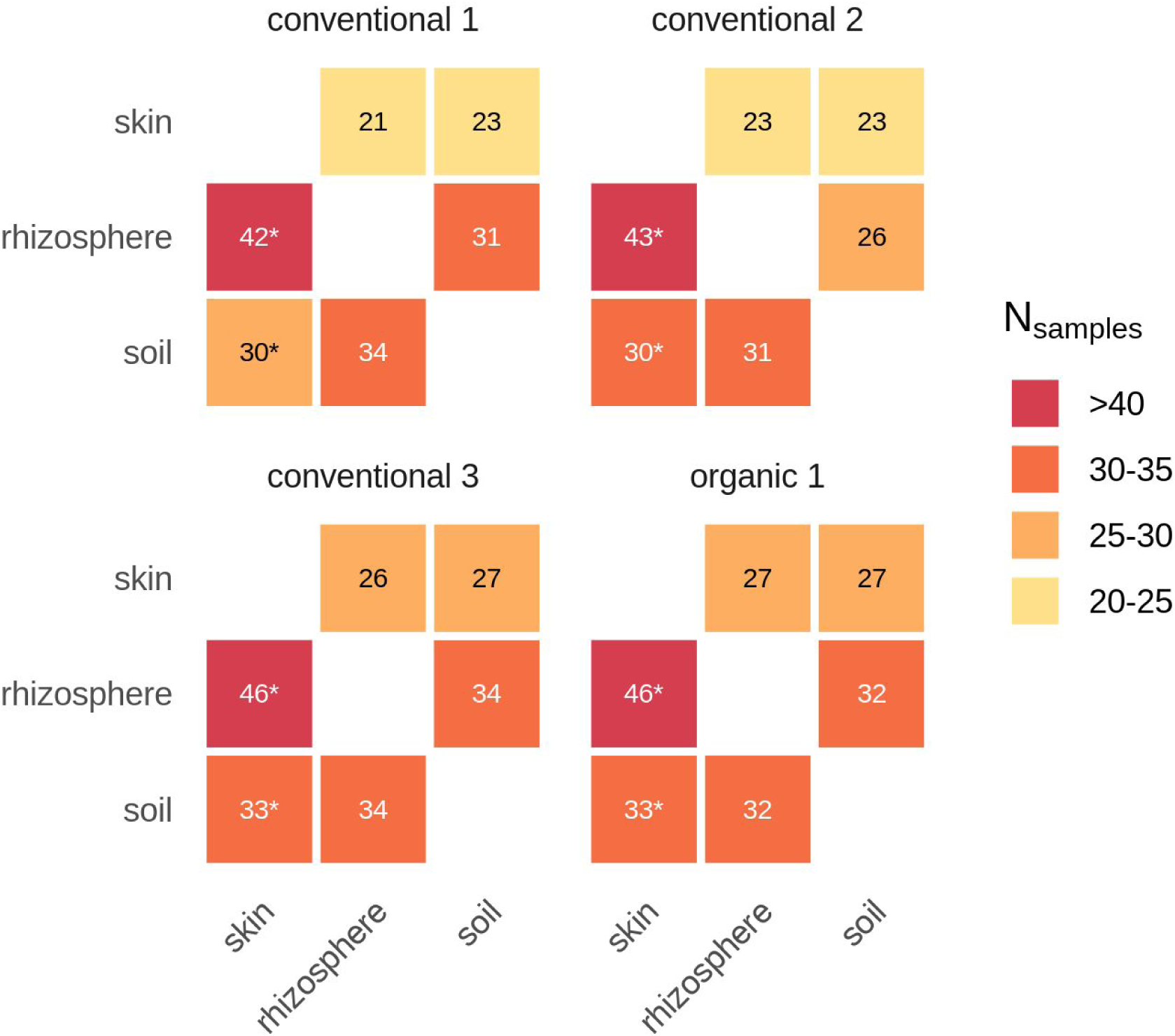
Nesting of compartment ASV membership. N_samples_ for all compartment pairwise comparisons within a given field. Presence/absence vectors of ASVs were ordered such that nesting of one compartment by another could be measured (see Methods). In this figure, cell values show the nesting of the compartment indicated in a column by the compartment indicated in the row. For example, the **bottom left cell** of each field location panel shows the nesting of sweetpotato **skin** by **soil**. For further information on field locations see Table 1. Asterisks indicate N_samples_ values with corresponding p-values less than 0.05.

**Supplementary Figure 7.**
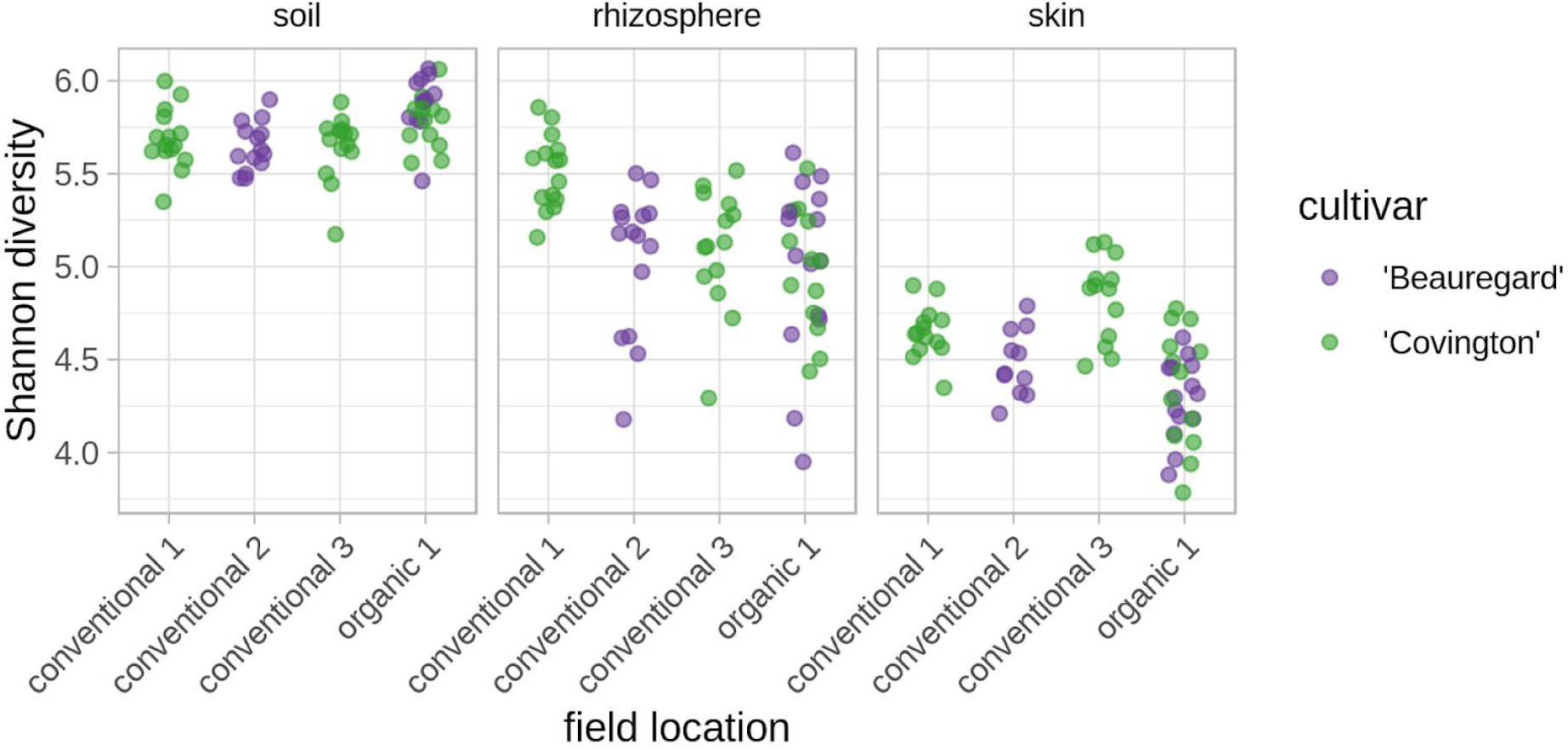
Shannon diversity of samples. Shannon diversity based on ASV content of all samples grouped by compartment and field location. Each point represents a sample and color denotes the sweetpotato cultivar.

**Supplementary Figure 8.**
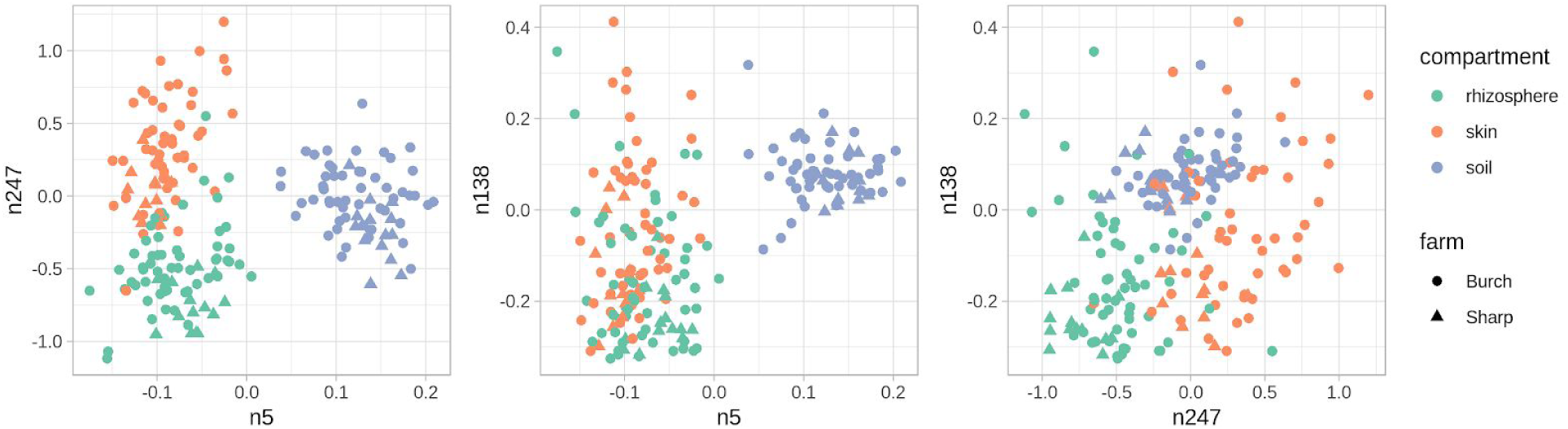
Phylogenetic ILR balance values for balances that separated compartments at both farms. Phylogenetic ILR values for the three balances that separate samples by compartment in both Burch Farm and Sharp Farm. Balance phylogenetic positions can be found in Figure 3, and Supplementary Figure 1. Balance ‘n5’ consists of a clade with consensus taxonomic affiliation of *Acidobacteria* (numerator clade) and a clade with consensus taxonomic affiliation of *Bacteria* (denominator clade). Balance ‘n138’ consists of a clade with consensus taxonomic affiliation of *Proteobacteria* (numerator clade) and a clade with consensus taxonomic affiliation of *Alphaproteobacteria* (denominator clade). Balance ‘n247’ consists of a clade with consensus taxonomic affiliation of *Planctomycetaceae* (numerator clade) and a clade with consensus taxonomic affiliation of *Bacteria* (denominator clade).

**Supplementary Figure 9.**
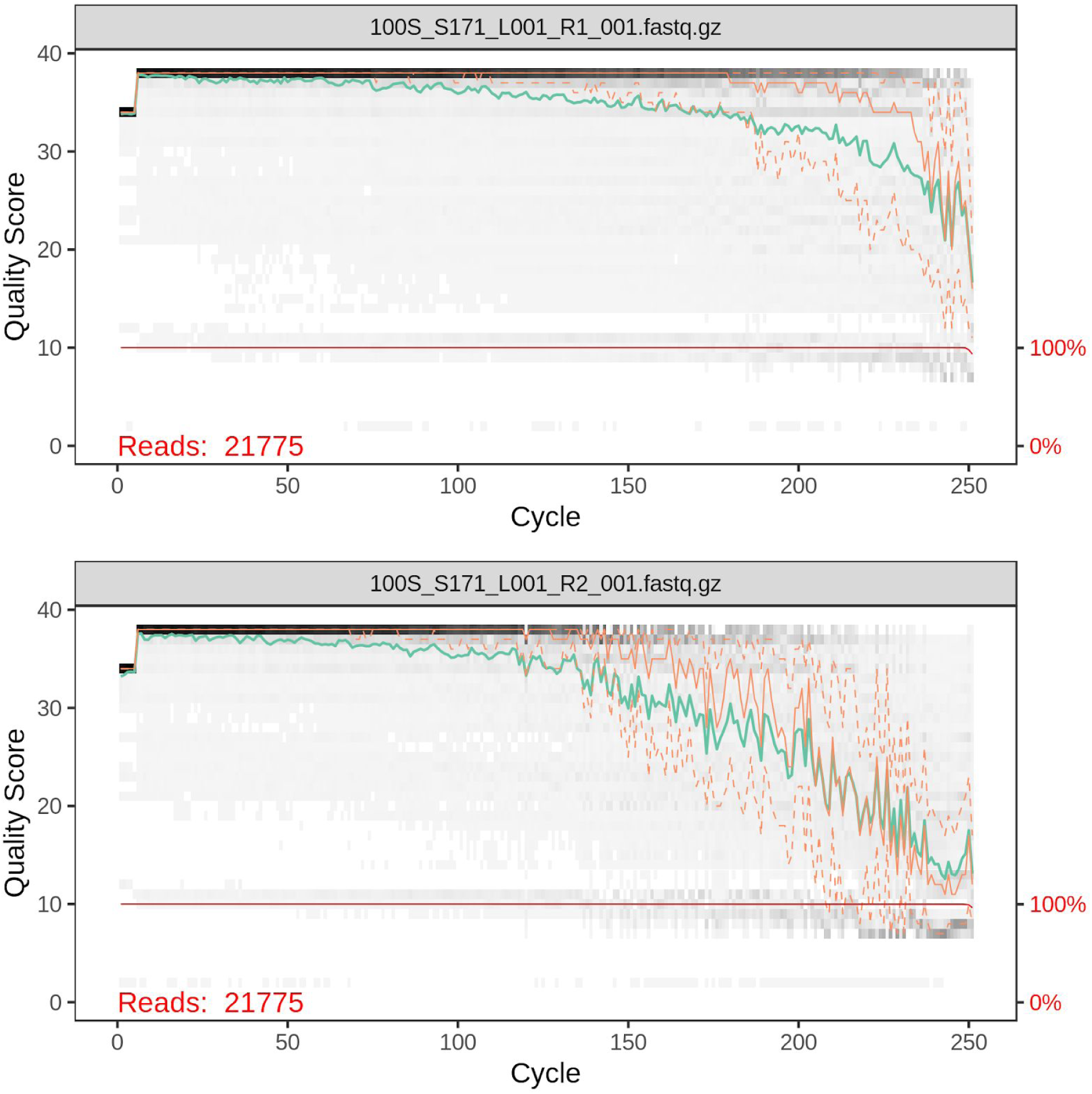
Quality score profile of DNA sequence reads. Example DNA-sequencing read quality score profiles for one sample. Darker cells denote greater frequency of base calls at the given quality score and cycle. The green line shows the mean quality score, solid orange is the median, and dashed orange lines indicate the 25th and 75th quality score quartiles. The red line shows the percentage of reads that extend to the indicated position.

**Supplementary Figure 10.**
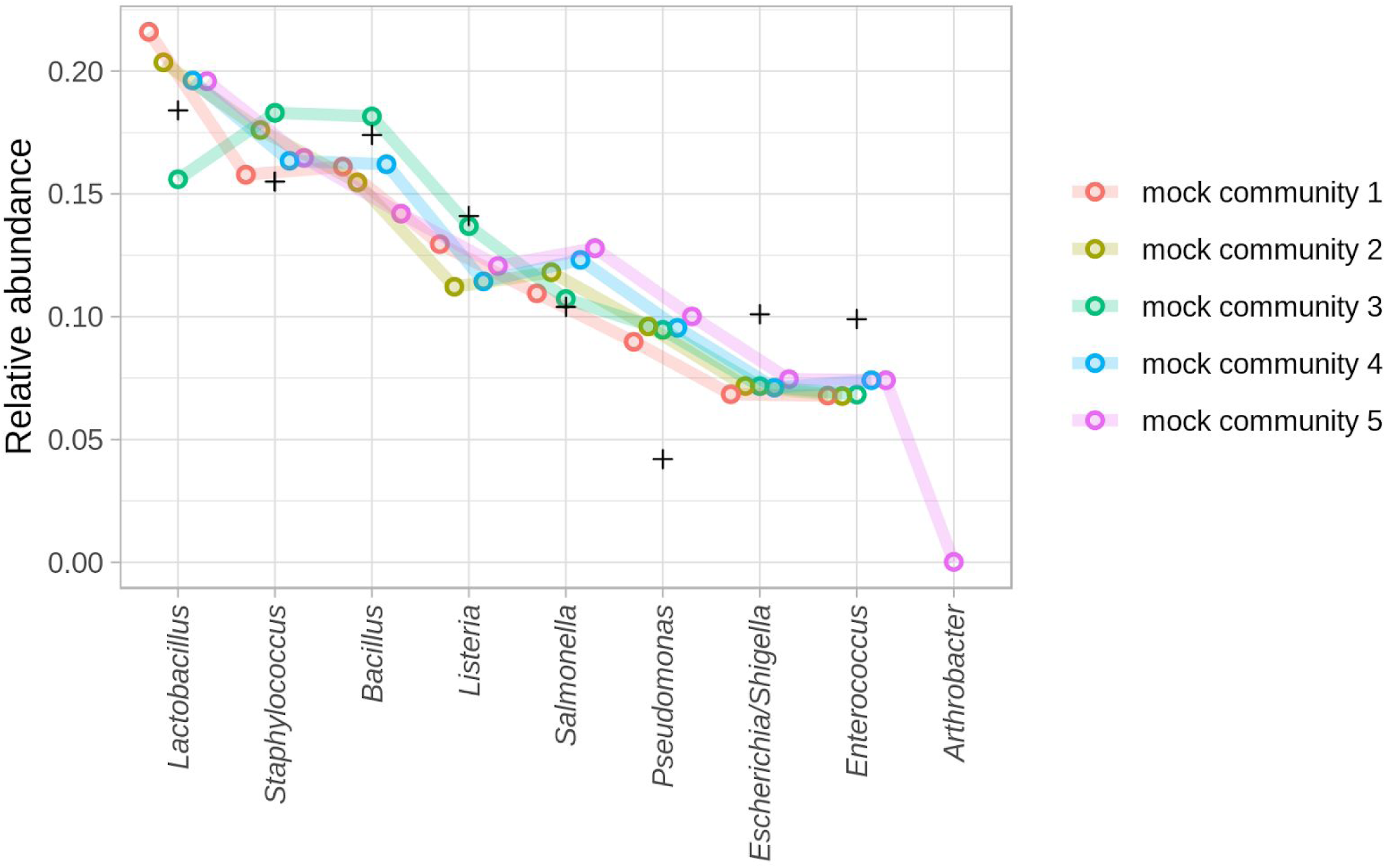
Mock community member relative abundances. Relative abundance of mock community members for each of five mock community samples. ‘+’ symbols show the manufacturer’s theoretical relative abundance estimate for each mock community member (ZymoBIOMICS, D6305; Irvine, CA).

**Supplementary Figure 11.**
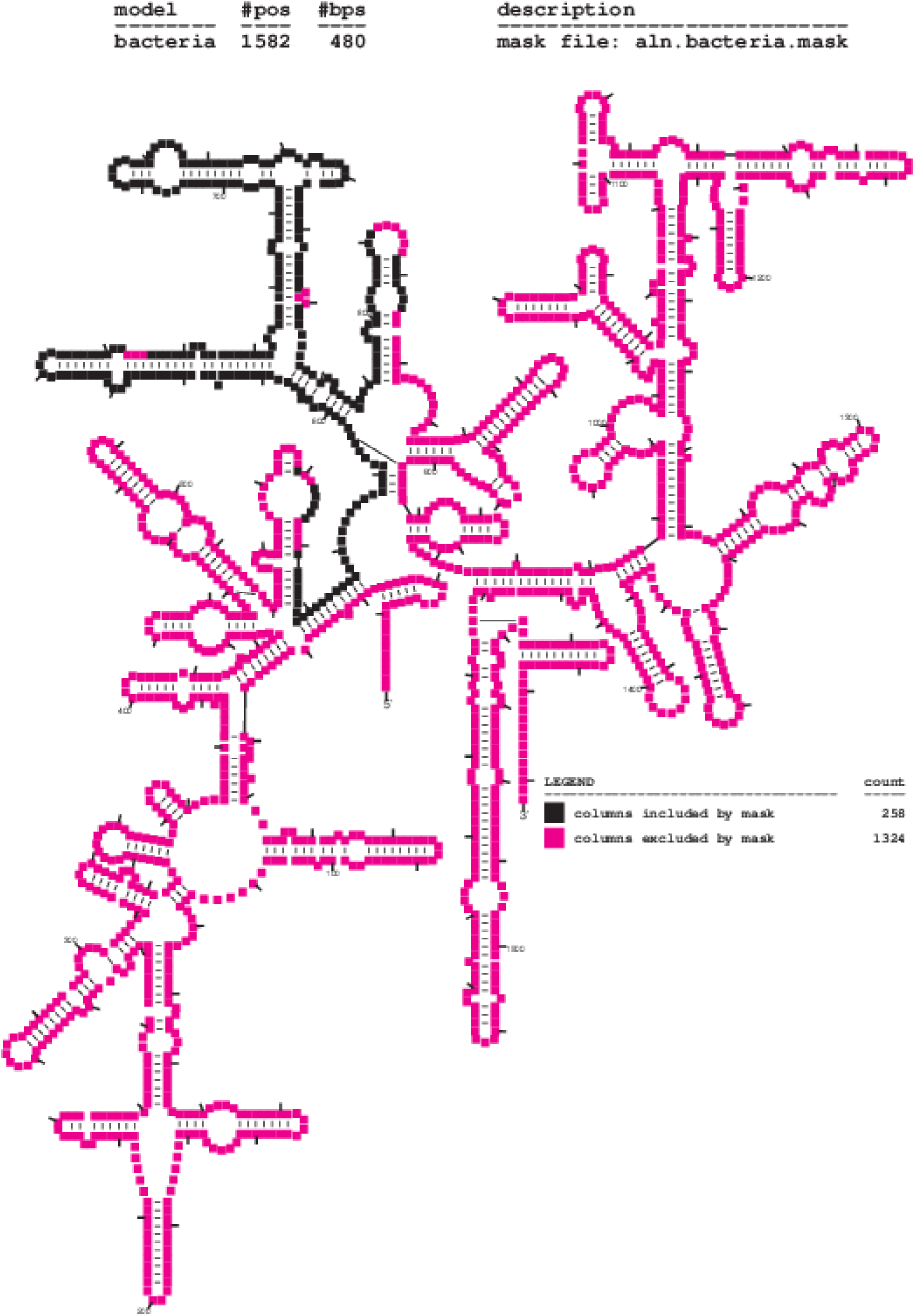
Alignment positions for phylogenetic reconstruction. Secondary structure diagram of the SSU rRNA gene with the V4 hypervariable positions we used for phylogenetic reconstruction highlighted in black (Nawrocki 2009).

